# CDX2 dose-dependently influences the gene regulatory network underlying human extraembryonic mesoderm development

**DOI:** 10.1101/2024.01.25.577277

**Authors:** Emily A. Bulger, Todd C. McDevitt, Benoit G. Bruneau

**Author notes:** Corresponding Author: Benoit G. Bruneau.

## Abstract

Proper regulation of gene dosage is critical for the development of the early embryo and the extraembryonic tissues that support it. Specifically, loss of *Cdx2 in vivo* leads to stunted development of the allantois, an extraembryonic mesoderm-derived structure critical for nutrient delivery and waste removal in the early embryo. In this study, we investigate how CDX2 dose-dependently influences the gene regulatory network underlying extraembryonic mesoderm development. We generate an allelic series for *CDX2* in human induced pluripotent stem cells (hiPSCs) consisting of WT, heterozygous, and homozygous null *CDX2* genotypes, differentiate these cells in a 2D gastruloid model, and subject these cells to multiomic single nucleus RNA and ATAC sequencing. We identify several genes that CDX2 dose-dependently regulate cytoskeletal integrity and adhesiveness in the extraembryonic mesoderm population, including regulators of the VEGF, canonical WNT, and non-canonical WNT signaling pathways. Despite these dose-dependent gene expression patterns, snATAC-seq reveals that heterozygous CDX2 expression is capable of inducing a WT-like chromatin accessibility profile, suggesting accessibility is not sufficient to drive gene expression when the CDX2 dosage is reduced. Finally, because the loss of CDX2 or TBXT phenocopy one another *in vivo*, we compare differentially expressed genes in our CDX2 knock-out model to those from TBXT knock-out hiPSCs differentiated in an analogous experiment. This comparison identifies several communally misregulated genes that are critical for cytoskeletal integrity and tissue permeability, including *ANK3* and *ANGPT1*. Together, these results clarify how CDX2 dose-dependently regulates gene expression in the extraembryonic mesoderm and suggest these genes may underlie the defects in vascular development and allantoic elongation seen in the absence or reduction of CDX2 *in vivo*.

**Summary Statement:** Using 2D human gastruloids, CDX2 is shown to dose-dependently influence genes related to tissue permeability, cell-cell adhesions, and cytoskeletal architecture during extraembryonic mesoderm development.

## INTRODUCTION

Throughout embryonic development, proper regulation of gene dosage is necessary for the morphogenesis of the embryo and the tissues that support it. One transcription factor with critical roles in both extraembryonic and embryonic tissue development is CDX2, a homeodomain-containing protein that is critical for placental development and later the axial elongation of the tailbud (Beck *et al*., 1995; van den Akker *et al*., 2002; Chawengsaksophak *et al*., 2004; Strumpf *et al*., 2005; Savory *et al*., 2009; Foley *et al*., 2019). CDX2 is first expressed during the blastocyst stage of embryogenesis during the specification of the trophectoderm, where it is required for proper implantation (Chawengsaksophak *et al*., 1997; Niwa *et al*., 2005). As development progresses toward gastrulation, CDX2 expression becomes localized to the posterior primitive streak where CDX2+ cells give rise to the extraembryonic mesoderm. The extraembryonic mesoderm contributes to the formation of the heavily vascularized allantois, enables the fusion of the allantois with the chorion in a process known as chorioallantoic fusion, and contributes to vasculogenesis of the yolk sac mesoderm. These structures are critical for nutrition, gas exchange, and waste removal in the early embryo, and their failure to develop properly leads to asphyxiation, nutrient deprivation, and premature death. As such, epiblast-specific *Cdx2* mutant mice have severely underdeveloped allantoic buds and these buds do not fuse with the chorion, preventing the formation of a functional chorioallantoic placenta (Chawengsaksophak *et al*., 2004; BrookelBisschop *et al*., 2017; Foley and Lohnes, 2022). Yolk sac vasculogenesis is also impaired in these mutants, preventing normal circulation. Despite thorough documentation of the physical manifestations induced by impaired *Cdx2* expression, the specific molecular role of CDX2 in regulating the morphogenesis of these extraembryonic structures is not fully understood, nor has it been explored in a human system.

Interestingly, the loss of *Cdx2* in mice phenocopies the loss of several genes involved in the canonical WNT signaling pathway, including *Tcf/Lef*, *Wnt3*, and *Tbxt* (Rashbass *et al*., 1991; Galceran *et al*., 1999; Rossant and Cross, 2001; Inman and Downs, 2006). Much like *Cdx2* mutant embryos, *Tbxt* mutant embryos are unable to survive past approximately E10 due to impaired allantois formation. Cells in the allantoic core where *Tbxt* would normally be expressed are significantly reduced in *Tbxt* mutants, however, cells of the outer mesothelium appear intact and these cells remain seemingly competent to adhere to the chorion (Inman and Downs, 2006). These observations suggest that *Tbxt* is necessary for the allantois to extend long enough to reach the chorion but is not necessarily required for chorioallantoic fusion itself. In addition, mice that are heterozygous for a functional *Tbxt* allele have an intermediate phenotype to WT or *Tbxt*-null mice, displaying variable allantois and blood island development and delayed vasculogenesis. Even so, allantoic growth in these heterozygotes is usually sufficient to allow for eventual chorioallantoic fusion, enabling pups to survive to term. Whether *Cdx2* heterozygosity also influences allantoic growth and chorioallantoic fusion is not yet known.

Acquisition of extraembryonic mesoderm identity is intimately tied to gastrulation, as this tissue emerges from the posterior primitive streak because of a network consisting of BMP, WNT, and NODAL signaling (Arnold and Robertson, 2009). *In vitro,* 48 hours of BMP4 exposure can induce this network in 2D cell colonies (“2D gastruloids”) in a way that reproducibly generates concentric rings of epiblast-like cells, embryonic mesoderm-like cells, endoderm-like cells, and extraembryonic mesoderm-like cells (Warmflash *et al*., 2014; Minn *et al*., 2020, 2021). This model allows us to investigate both how specific genes active during early gastrulation augment cell identity and how changes in adjacent tissues influence cell-cell communication and the gene regulatory networks underlying lineage emergence.

In this study, we employ this 2D gastruloid model and subsequent multiomic single nucleus RNA sequencing (snRNA-seq) and Assay for Transposase Accessible Chromatin sequencing (snATAC-seq) to identify how CDX2 regulates the proper morphogenesis of extraembryonic mesoderm and the extent to which this regulation is controlled in a dose-dependent manner during early gastrulation. We demonstrate that varying CDX2 dose at this stage of early development directly influences genes involved in cell-cell adhesions, extracellular matrix integrity, cytoskeletal architecture, and HOX expression, in addition to influencing key regulators of extraembryonic mesoderm fate such as the WNT and GATA families of factors. We then compare this dataset to an analogous study looking at the impact of TBXT dose on extraembryonic mesoderm development (Bulger *et al*., 2023). This comparison allows us to further isolate several genes with shared misregulation that are involved in cytoskeletal integrity and tissue permeability, including *ANK3* and *ANGPT1,* which are both involved in VEGF-directed blood vessel maturation. Taken together, this study suggests that CDX2 activates gene regulatory networks associated with impaired allantois formation and vasculogenesis in a dose-dependent manner, including but not limited to genes involved in cell adhesions, motility, and membrane permeability.

## RESULTS

### Generation of a hiPSC CDX2 allelic series and 2D gastruloids

To explore the effect of CDX2 dose on the development of the extraembryonic mesoderm population, we engineered a WTC11-LMNB1-GFP-derived allelic series consisting of CDX2^+/+^ (WT), CDX2^+/-^ (CDX2-Het), and CDX2^-/-^ (CDX2-KO) human induced pluripotent stem cell lines (hiPSCs). These mutants were generated by targeting the first exon of CDX2 with CRISPR/Cas9, generating a premature stop codon on one or two alleles, respectively, as confirmed by Sanger sequencing (Fig. 1A, S1, Table S1, methods). We conducted a western blot for CDX2 in a sparsely seeded monolayer of cells exposed to BMP4 for 48 hours and this revealed the expected stepwise decrease in CDX2 expression across the allelic series, with the highest protein levels in the WT, intermediate levels in CDX2-Het, and no detectable protein in CDX2-KO cells (Fig. 1B). The complete absence of CDX2 protein in CDX2-KO cells was further confirmed through immunofluorescence (IF) (Fig. 1D-E).

**Figure 1:**
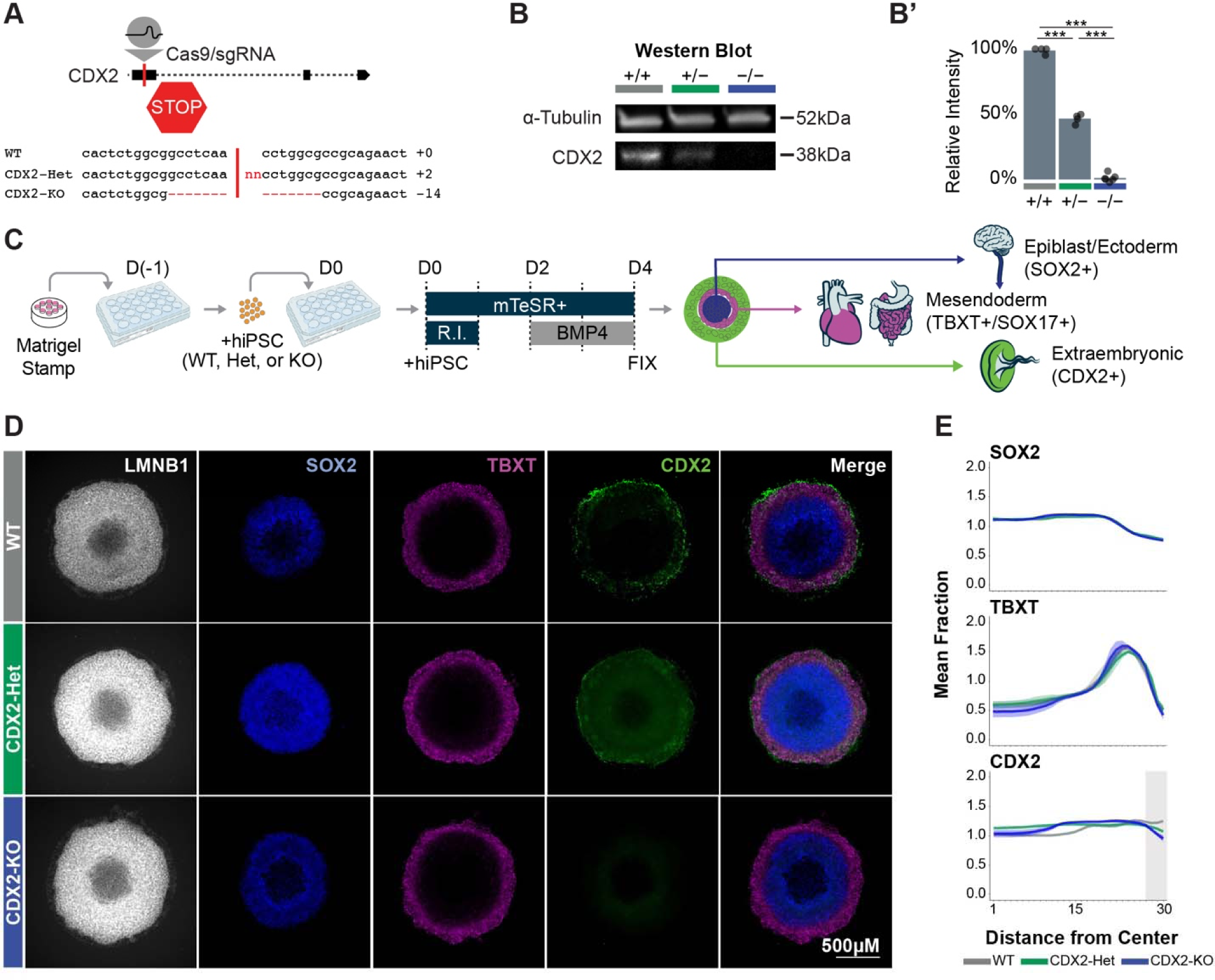
Generation and validation of the CDX2 allelic series. (**A**) Schematic of CDX2 locus and sgRNA target in the first exon. Indel generated in CDX2-Het (+2bp) or CDX2-KO (−14bp). (**B**) Western blot and (**B’**) quantification CDX2 western blot signal intensity in WT, CDX2-Het, and CDX2-KO after 48hr BMP4 exposure. n=4 replicates/genotype. (**C**) Schematic of differentiation protocol. (**D**) Immunofluorescence for SOX2, TBXT, and CDX2 in WT, CDX2-Het, and CDX2-KO 2D gastruloids. Nuclei labeled with LMNB1. (**E**) Quantification of the mean fraction of fluorescence intensity across gastruloids of each genotype. 1 = center of gastruloid, 30 = outer gastruloid. n = 3-7 gastruloids/genotype.

### Lineage emergence is minimally altered in 2D gastruloids of varying CDX2 dose

To investigate how CDX2 dose influences extraembryonic mesoderm specification and morphogenesis, we differentiated the allelic series into 2D gastruloids via 48 hours of BMP4 exposure (Warmflash *et al*., 2014; Minn *et al*., 2020) (Fig. 1C, methods). We then conducted two bioreplicates of multiomic single-nucleus RNA-sequencing (snRNA-seq) and single-nucleus Assay for Transposase-Accessible chromatin (snATAC-seq) (Buenrostro *et al*., 2015) on gastruloids of each genotype after 48 hours of BMP4 treatment. Through this in-depth sequencing approach, we sought to precisely define how CDX2 dose influences cell identity and the expression of morphogenesis regulators during early gastrulation. In addition, CDX2 has been shown to sustain newly accessible chromatin regions in mature tissues (Verzi *et al*., 2010, 2013; Neijts *et al*., 2017; Saxena *et al*., 2017; Kumar *et al*., 2019) and also to interact with the Brg1 subunit of the switch-sucrose non-fermentable (SWI-SNF) chromatin remodeling complex (Yamamichi *et al*., 2009; Nguyen *et al*., 2017). We were interested in the extent to which CDX2 can dose-dependently influence chromatin accessibility in this nascent extraembryonic tissue.

After computationally pooling cells of all three genotypes and conducting dimensionality reduction and clustering of snRNA-seq data using the R package, Seurat (Satija *et al*., 2015), our analysis yielded ten distinct clusters reflecting various populations expected in the gastrulating embryo. This included three extraembryonic cell populations (C1–C3; “Extraembryonic-1–3”, epiblast-like cells (C4; “Epiblast”), three primitive streak-like cell populations (C5–C7; “PS-1–3“), nascent mesoderm-like cells (C8; “Mesoderm”), nascent endoderm-like cells (C9; “Endoderm”), and primordial germ cell-like cells (C10; “PGCLC”) (Fig. 2A-C, S2-3). The proportion of cells of each genotype assigned to each cluster was largely consistent, suggesting that the acquisition of cell identity during early gastrulation is not significantly affected by the loss of CDX2 (Fig. 2A’).

**Figure 2:**
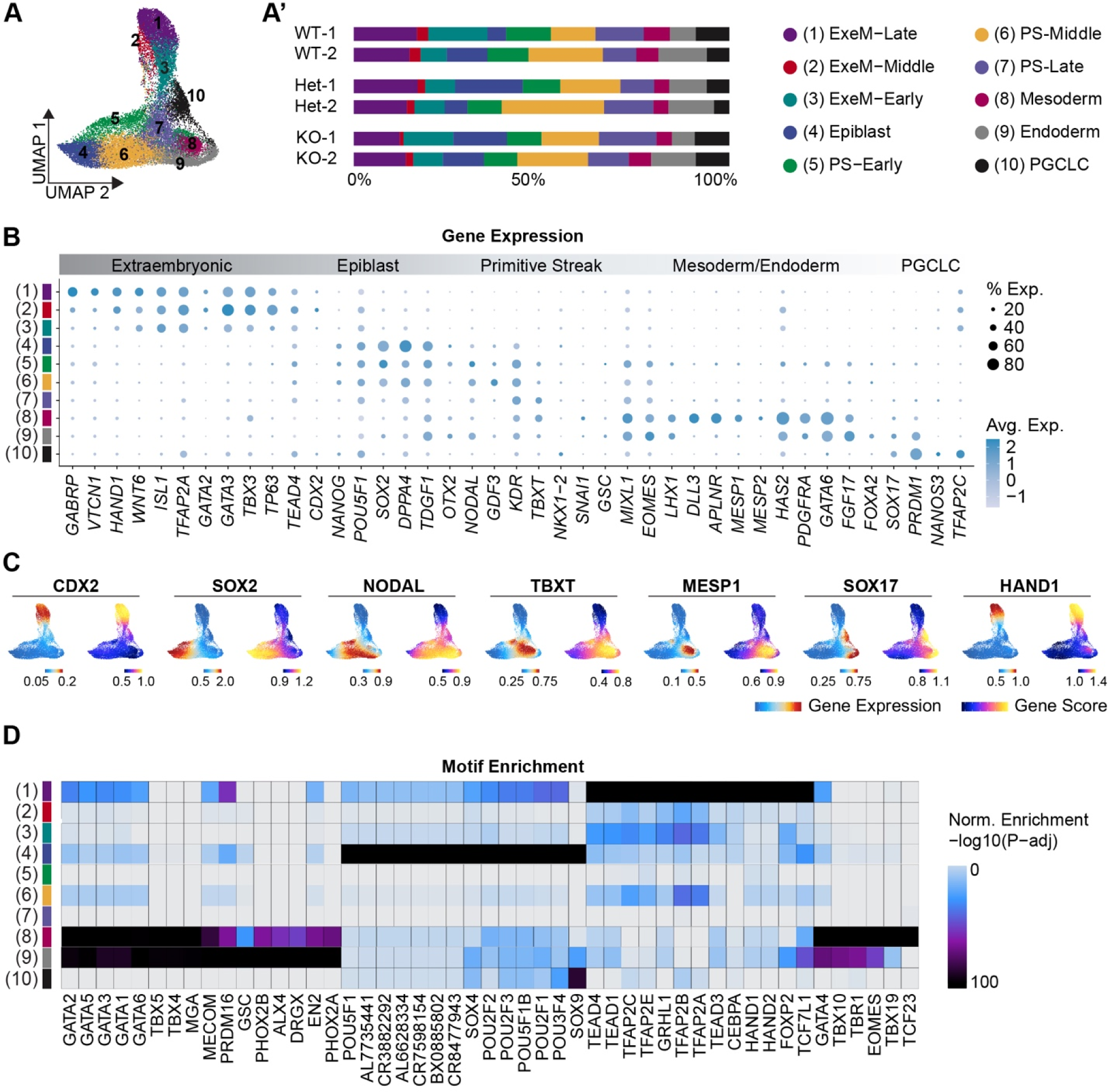
Lineage emergence is minimally altered in 2D gastruloids of varying CDX2 dose. (**A**) UMAP reflecting 10 clusters comprised of WT, CDX2-Het, and CDX2-KO cells from 2D gastruloids. (**A’**) Proportion of cells of each sample in each cluster. (**B**) Dot plot depicting key lineage markers across all 10 clusters. (**C**) Feature Plots reflecting the gene expression (snRNA-seq) or gene score (snATAC-seq) of key lineage markers. (**D**) Heatmap of the top motifs enriched in each cluster (FDR <= 0.1 & abs(Log2FC) >= 0.5, maximum motifs per cluster plotted = 20).

The extraembryonic cell type that emerges in 2D gastruloids has previously been shown to express genes that are shared between extraembryonic mesoderm/amnion and trophectoderm, including *CDX2, GATA3, TFAP2A, HAND1,* and *WNT6* (Bernardo *et al*., 2011; Warmflash *et al*., 2014; Ma *et al*., 2019; Niu *et al*., 2019; Zheng *et al*., 2019; Minn *et al*., 2020). To distinguish between these possible cell types and identify differences between the three extraembryonic clusters, we evaluated these clusters for key markers of trophectoderm, late-amnion, and early-amnion lineages. This analysis revealed a bias in all three clusters toward late-amnion identity (*GAPBRP, HEY1, HAND1, VTCN1, TPM1, IGFBP3, ANKS1A*) relative to embryonic clusters, in agreement with published findings that BMP4 drives primed hiPSCs toward an amnion-like fate (Rostovskaya *et al*., 2022) and reflecting an extraembryonic mesoderm rather than trophectoderm origin (Fig. S5). This late-amnion expression pattern was the highest in extraembryonic-1 and the lowest in extraembryonic-3, suggesting extraembryonic-1 is a relatively more differentiated extraembryonic mesoderm while extraembryonic-3 remains more nascent. Upon analysis using the R package ArchR (Granja *et al*., 2021), the companion snATAC-seq dataset revealed that the accessible regions of chromatin in cells within these extraembryonic clusters, whose identity information was imported from our snRNA-seq analysis based on matching cell barcodes (methods), are enriched for motifs for the TEAD, TFAP, and HAND families of transcription factors, which are critical regulators of extraembryonic mesoderm fate (Fig. 2D, S3-4). Again, we observe a stepwise enrichment pattern where these motifs are most highly enriched in extraembryonic-1 and have variable enrichment in extraembryonic-2 and extraembryonic-3. We therefore designated clusters 1-3 as extraembryonic-mesoderm-like late (“ExeM-Late”), middle (“ExeM-Middle”), and early (“ExeM-Early”), respectively. This annotation coincided with the trajectory analysis which ordered cells in pseudo time from ExeM-Early to ExeM-Late (Fig. S5B)

The remaining seven clusters were identified as outlined in Bulger et. al., 2023, which shares a WT dataset with the current analysis (Fig. 2B-C). Briefly, cluster 4 exhibits canonical hallmarks of epiblast cell fate including gene expression of *POU5F1*, *SOX2*, and *NANOG.* The three primitive streak clusters share canonical elements of the PS gene signature including *TBXT, MIXL1,* and *EOMES,* and are differentiated from one another by the progressive downregulation of epiblast markers such as *SOX2, TDGF1*, and *NODAL*. Based on this observation, we designated “PS-1” (C5) as “PS-Early”, “PS-2” (C6) as “PS-Middle”, and “PS-3” (C7) as “PS-Late.” Clusters 8 and 9 share a mesendoderm-like signature including enriched GATA and TBX family motifs and are distinguished by a mesoderm-like gene expression signature in cluster 8 (*MESP1, MESP2, ALPNR, TBXT*) and an endoderm-like gene expression pattern in cluster 9 (*FOXA2, SOX17, PRDM1*) (Fig. 2B-D). Lastly, cluster 10 reflects PGCLC identity, as evidenced by the coexpression of *SOX17, PRDM1, NANOS3*, and *TFAP2C*. The SOX9 motif was also enriched in the accessible regions that distinguish cells of this cluster, in agreement with its role as a marker of PGCLC-derived Sertoli cells later in development (Hayashi *et al*., 2018) (Fig. 2D).

### CDX2 primarily influences the GRN underlying extraembryonic mesoderm identity

To understand how CDX2 influences early embryonic morphogenesis, we next isolated each cluster and compared the number of differentially expressed genes (DEGs) between WT vs. CDX2-Het, WT vs. CDX2-KO, or CDX2-Het vs. CDX2-KO. As expected, the ExeM-Late cluster yielded both the highest CDX2 expression in WT cells (Fig. 3A) and the largest number of DEGs between WT and CDX2-KO relative to the other clusters (Table S2). Based on these changes in expression and because of the biological relevance of this cluster to extraembryonic mesoderm development, we focused on this cluster for subsequent analyses (Fig. 3B).

**Figure 3:**
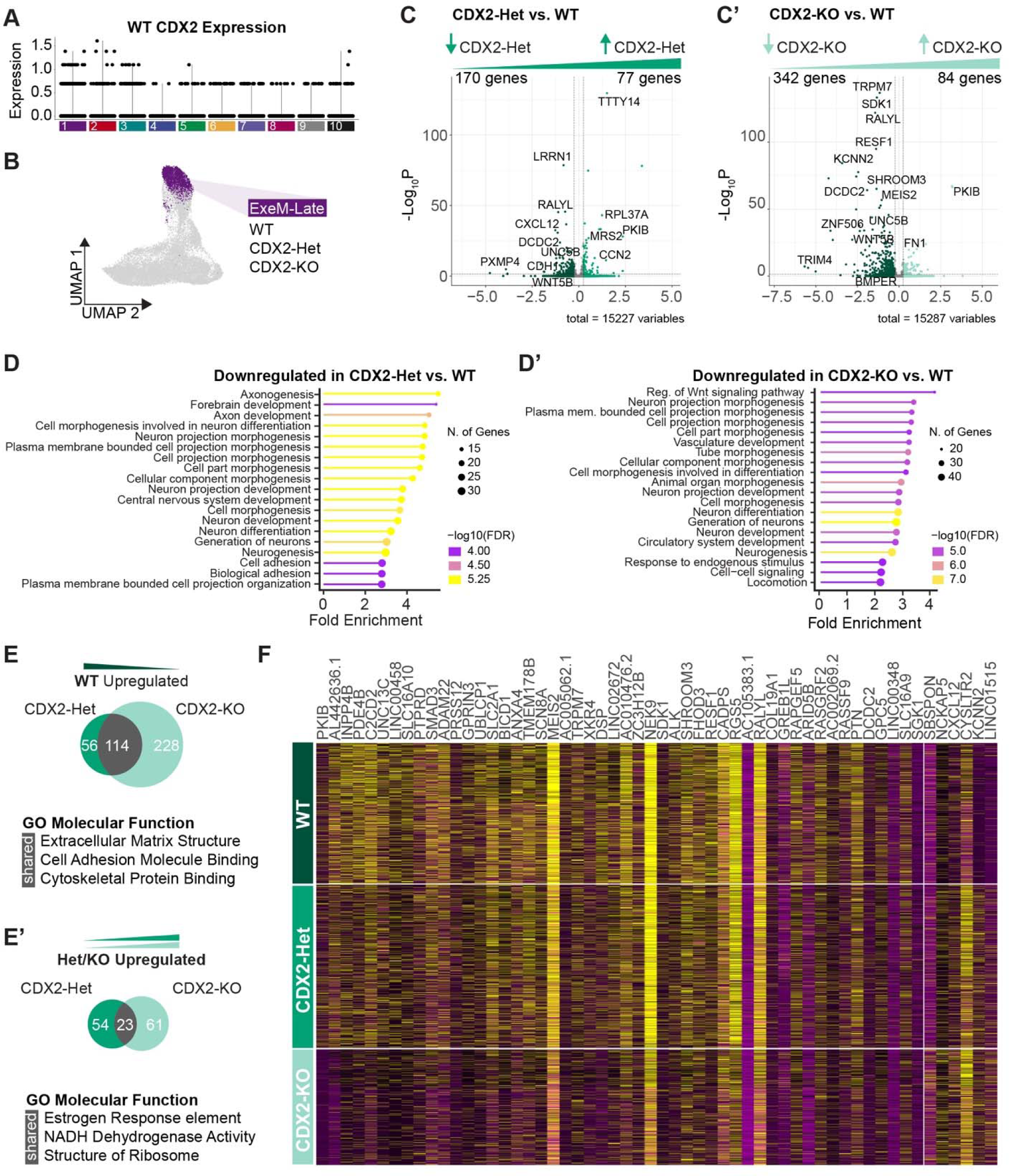
CDX2 dose-dependently influences extraembryonic mesoderm gene expression. **(A)** Violin plot of CDX2 expression across clusters in WT cells. **(B**) ExeM-Late cluster isolated for genotype-specific analyses. (**C**) Volcano plot of differentially expressed genes comparing WT and CDX2-Het or (**C’**) WT and CDX2-KO (abs(Log2FC) > 0.25, p-adj < 0.05) within the ExeM-Late cluster. (**D**) GO biological process enrichment for genes downregulated in CDX2-Het relative to WT or (**D’**) CDX2-KO relative to WT (abs(Log2FC) > 0.25, p-adj < 0.05, ShinyGO FDR < 0.05) within the ExeM-Late cluster. (**E**) Venn diagram of genes upregulated or (**E’**) downregulated in WT relative to CDX2-Het (left) or CDX2-KO (right). Select results from GO molecular function enrichment for overlapping genes. (**F**) Heatmap of the top differentially expressed genes between WT and CDX2-KO (p-adj < 0.05, abs(Log2FC >1, pct cells > 20%) within the ExeM-Late cluster.

Comparing CDX2-Het to WT, we detected 170 genes downregulated and 77 genes upregulated in the CDX2-Het (Fig. 3C, Table S2). Downregulated genes included canonical WNT inhibitor *DCDC2*, cell-cell adhesion regulator *CDH1*, WNT ligand *FZD6*, and BMP inhibitor *BMPER,* while the upregulated genes largely consisted of ribosomal proteins. GO biological process enrichment for the genes downregulated in CDX2-Het revealed several pathways related to “cell projection morphogenesis” and “cell adhesion”, and also several neural-related pathways (Fig. 3D, Table S3). These neural pathways are largely driven by genes involved in axon guidance such as *FYN*, *UN5B*, and *SEMA3E*, so while they may reflect a neural bias in cell identity it is also probable that they reflect broader patterns in misregulated cell migration.

Comparing CDX2-KO to WT, we detected 342 genes downregulated and 84 genes upregulated in the CDX2-KO (Fig. 3C’, Table S2). Like in CDX2-Het, downregulated genes in CDX2-KO included several regulators of the WNT signaling pathway including *WNT5B*, *DCDC2*, and *FZD6*, as well as several genes related to cell-cell adhesion including *CDH1* (Fig. 3C’, Table S2). Genes involved in cell migration and axon guidance such as *GLI3*, *UNC5B*, *SEMA3E*, and *ISL1* were also identified. GO biological process enrichment revealed a 4-fold enrichment of genes involved in “canonical WNT signaling,” as well as enrichment for “vasculature development”, “circulatory system development”, “tube morphogenesis”, “cell-cell signaling”, and “locomotion” (Fig. 3D’, S7, Table S3) Taken together, the differentially expressed genes that are seen in both CDX2-Het and CDX2-KO vs. WT correspond to several properties of the developing allantois and other extraembryonic-mesoderm-derived structures including cell projection morphogenesis, adhesions, and WNT signaling, and potentially underly the impaired vasculogenesis phenotype observed in mouse models.

To clarify the extent to which these pathways are dependent on CDX2 dosage, we next determined the number of overlapping downregulated genes from both the CDX2-Het vs. WT and the CDX2-KO vs. WT comparisons. Of the 170 genes downregulated in CDX2-Het and the 342 genes downregulated in CDX2-KO, we found that 114 genes were conserved including *SMAD3*, *MEIS2*, and *DCDC2* (Fig. 3E-F, Table S2) GO Molecular Function analysis of these 114 genes revealed significant enrichment for “extracellular matrix structure”, “cell adhesion molecule binding”, and “cytoskeletal protein binding” (Fig. 3E, S7, Table S3). GO biological process enrichment for “vasculature development” remained enriched by approximately 3-fold (FDR = 0.03), and of the 29 genes driving “vasculature development” enrichment in CDX2-KO, 10 remained differentially expressed in CDX2-Het (*ISL1, ATP2B4, GLI3, ENG, UNC5B, GPLD1, RTN4, COL4A2, HEY1, SEMA3E*) (Table S2). In contrast, there were 23 genes upregulated in both CDX2-Het and CDX2-KO, and the GO enrichment terms for these genes included “estrogen response element”, “NADH dehydrogenase activity” and “structure of ribosome.” However, it is difficult to evaluate the biological significance of these terms due to the low number of overlapping genes (Fig. 3E’, Table S3). These results demonstrate that CDX2 dose-dependently influences both physical cell structure and the gene regulatory network involved in the development of extraembryonic tissue and largely acts as a transcriptional activator.

### snATAC-seq reveals loss of accessibility in regions with CDX2 motifs

To better understand how CDX2 dosage influences the gene regulatory network underlying extraembryonic-mesoderm development, we next turned to the paired snATAC-seq dataset generated from the same nuclei as our snRNA-seq dataset. As for snRNA-seq, we subset the ExeM-Late cluster and identified regions with differential chromatin accessibility (differentially accessible regions; DARs) between either WT and CDX2-Het or WT and CDX2-KO populations.

Through this analysis, we identified very few DARs between CDX2-Het and WT in the Exe-Late cluster (13/174,048 DARs with increased accessibility in CDX2-Het and 12/174,048 DARs with increased accessibility in WT) (Table S2). In contrast, comparing CDX2-KO to WT in the same cluster revealed many more DARs, as we detected 29/174,048 DARs with increased accessibility in CDX2-KO and 232/174,048 DARs with increased accessibility in WT (Fig. 4A, Table S2). The relatively high number of reduced-accessibility peaks detected in the CDX2-KO compared to the WT condition suggests that CDX2 is either directly or indirectly responsible for making specific regions of chromatin more accessible in the extraembryonic mesoderm population, in agreement with studies showing CDX2 can serve as a pioneer factor (Zaret and Mango, 2016; Neijts *et al*., 2017). Additionally, the lack of DARs detected in the CDX2-Het vs. WT comparison indicates that an intermediate level of CDX2 is sufficient to induce a WT-like chromatin accessibility profile (Fig. 4B).

**Figure 4:**
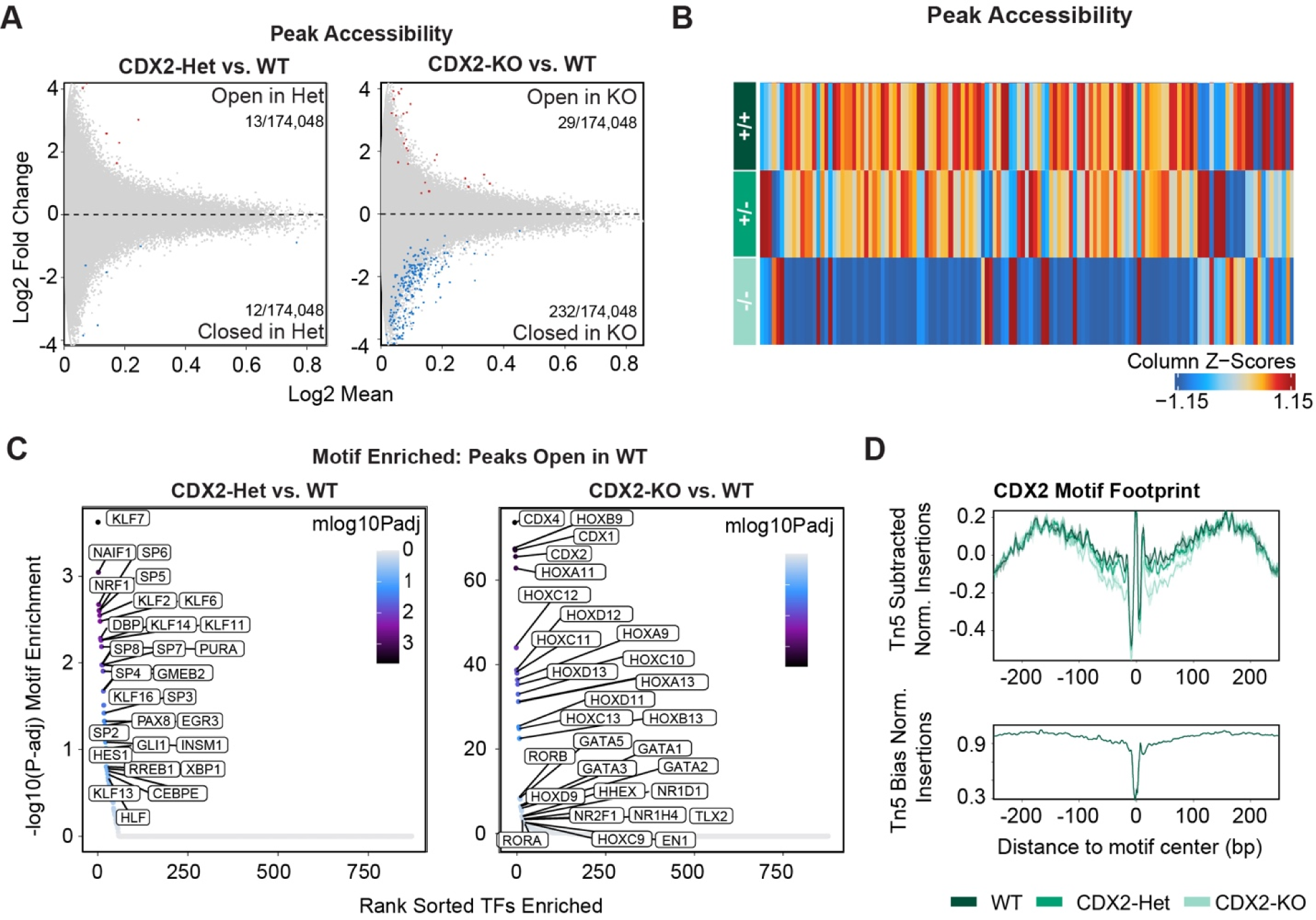
CDX2 expression influences chromatin organization at regions containing CDX motifs. **(A)** Differential peak accessibility comparing CDX2-Het and WT (left) or CDX2-KO and WT (right) within the ExeM-Late cluster (FDR < 0.1 & abs(Log2FC) > 0.5). Red = regions more accessible in mutant, blue = regions more accessible in WT. (**B**) Heatmap of Differentially accessible peaks across WT, CDX2-Het, and CDX2-KO within the ExeM-Late cluster (**C**) Enriched motifs detected in peaks uniquely accessible in WT relative to CDX2-Het (left) and CDX2-KO (right). (**D**) Footprint for the CDX2 motif across WT (dark green), CDX2-Het (middle green), and CDX2-KO (light green).

To better understand the composition of DARs that are uniquely accessible in each CDX2 condition, we next conducted motif enrichment to look for motifs that are overrepresented in these regions. Motifs enriched in peaks that are differentially accessible between WT and CDX2-Het or WT and CDX2-KO include proliferation regulators *KLF7*, *SP6*, and *GATA1*, but these motifs are enriched at a relatively low level (−log10(p-adj) ∼ 3) and may be driven by the low number of DARs found in these conditions (Fig. 4C, S8). In contrast, the top motifs in DARs that are more accessible in WT but less accessible in CDX2-KO exhibited significantly higher enrichment scores and consisted almost exclusively of homeobox genes, including *CDX2*, *CDX4*, and posterior *HOX* genes, and also *GATA* factors (−log10(p-adj) ∼60) (Fig. 4C). While this may reflect the established role of CDX2 in regulating HOX gene expression and associated chromatin dynamics (Amin *et al*., 2016; Neijts *et al*., 2017), CDX genes and posterior HOX genes share very similar binding motif (TTAT) that is distinct from anterior or central HOX genes (Ekker *et al*., 1994; Berger *et al*., 2008; Noyes *et al*., 2008; Mann, Lelli and Joshi, 2009; Bulajić *et al*., 2020). These hits therefore likely capture the redundancy in these motif annotations and highlight how CDX2 expression correlates with the accessibility of regions containing its binding motifs, suggesting it has the capability to remodel chromatin in regions where it is bound.

We next conducted transcription factor footprinting analysis to understand how TF binding at these CDX motifs is influenced by CDX2 dosage. As anticipated, we observe a lower footprint at the CDX2 motif for CDX2-KO reflective of reduced occupancy relative to WT (Fig. 4D). The footprint in the CDX2-Het was intermediate between WT and CDX2-KO, suggesting that the reduced gene expression correlates with reduced TF binding at CDX motifs. Because TF footprinting denotes average motif occupancy, it is unclear whether this is because fewer CDX TFs bind to a specific locus in favor of another with higher affinity or due to a global reduction in CDX TF binding across all loci containing CDX motifs. However, the lack of differentially accessible peaks observed in CDX2-Het vs. WT indicates that the intermediate CDX2 expression level and subsequent reduced occupancy of CDX motifs remains sufficient to induce a WT-like chromatin accessibility profile, despite the lower dose influencing gene expression levels for markers related to cell architecture and adhesions.

### CDX2 dose-dependently influences expression within the HOXB locus

Because CDX and posterior HOX motifs share a common binding motif that was identified as enriched in DARs, and because CDX2 influences HOX gene expression in the embryo proper(Neijts *et al*., 2017), we next asked how CDX2 dosage influences HOX gene expression patterns in the extraembryonic mesoderm. HOX clusters can be subdivided into three subsections containing paralogs A-D; the 3’ cluster containing HOX genes 1-4, the middle cluster containing HOX genes 5-9, and a 5’ cluster consisting of HOX genes 10-13 (Neijts *et al*., 2017), The 3’ cluster is thought to be activated by WNT signaling, while all paralogs within the middle cluster depend on CDX-transcription factors to become accessible. The 5’ cluster is activated much later in development by central HOX genes in a colinear fashion (Neijts *et al*., 2017). In agreement with previous work documenting an anterior homeotic shift in CDX2 mutant mice (van den Akker *et al*., 2002), we observed slightly elevated expression of the central HOX genes, specifically *HOXB3*-*HOXB7*, in WT relative to CDX2-KO (Fig. S6A). Most genes from *HOXA*, *HOXC*, and *HOXD* paralogs were not detected in our dataset, and this may be because the HOXB cluster typically slightly precedes the expression of other paralogs (Denans, Iimura and Pourquié, 2015). Alternatively, HOX paralogs have been shown to exert different roles in different tissues (Kachgal, Mace and Boudreau, 2012), and *HOXB3*, *HOXB5*, and *HOXB7* specifically have been shown to impact vasculogenesis (Miano *et al*., 1996; Myers, Charboneau and Boudreau, 2000; Wu *et al*., 2003). It is, therefore, possible that HOXB paralogs serve a unique role in early extraembryonic mesoderm development and subsequent placental vasculogenesis.

Like CDX2-KO, central HOX expression in CDX2-Het was also reduced relative to WT (Fig. S6A). This suggests that while intermediate CDX2 expression may be sufficient to induce changes in chromatin accessibility, a higher threshold of expression may be required to activate downstream HOX genes to WT levels. In contrast, the anterior HOX gene *HOXB2* was higher in CDX2-KO than both CDX2-Het and WT, likely because of the established role of more posterior HOX genes in suppressing the function of more anterior HOX genes (Chisaka and Capecchi, 1991; Lufkin *et al*., 1991, 1992; Duboule and Morata, 1994; Iimura and Pourquié, 2006; Denans, Iimura and Pourquié, 2015) (Fig. S6A). The posterior HOX genes, specifically *HOXB9*, *HOXA13*, and *HOXC13*, were detected in very few cells. Taken together, these results illustrate how reduced CDX2 dose-dependently limits the expression of downstream central HOX genes.

We next looked at snATAC-seq tracks between the three genotypes across the HOX paralogs. Through comparing these tracks to gene expression patterns, we were able to annotate “peak2gene links” highlighting specific peaks that correlate with changes in gene expression. Due to variability in gene expression, far more peak2gene links were detected in the HOXB cluster compared to HOXA, HOXC, and HOXD (Fig. S6B). Many of these peak2gene links correspond to genotype-specific chromatin organization, reflecting the ability of CDX2 to alter chromatin accessibility around HOX genes.

Overall, these results reflect the ability of CDX2 to augment HOX expression, including the downregulation of central HOX genes and upregulation of anterior and posterior HOX genes, in a dose-dependent and possibly paralog-specific manner.

### Cellchat reveals a dose-dependent role for CDX2 in regulating the non-canonical WNT signaling pathway

In the early extraembryonic mesoderm, paracrine and juxtracrine signals from a variety of tissues are required to orchestrate morphogenesis (Stewart, 1996; Downs *et al*., 2009). With this in mind, we utilized the R package CellChat (Jin *et al*., 2021) to identify patterns of ligand-receptor communication across clusters. We first investigated how CDX2 dosage influences broad patterns in pathway activation by isolating pathways with identifiable changes in “information flow”, which predicts patterns in cell-cell communication by quantifying the changes in signals between or within cell types.

This analysis revealed that the amount of information flow across most pathways is largely conserved across CDX2 genotypes, except the non-canonical WNT (ncWNT) signaling pathway which has reduced information flow in both CDX2-Het and CDX2-KO (Fig. 5A). We visualized the precise changes in predicted communication between and within clusters in circle plots, where line thickness correlates with the degree of predicted communication. This analysis revealed that the marked decrease in ncWNT communication within CDX2-Het and CDX2-KO was most clearly derived from changes in the three ExeM clusters (C1-C3), where both paracrine signaling (loops) and juxtracrine signaling (lines) are reduced (Fig. 5B). We quantified the relative contribution of the different ligands and receptor pairs that define the ncWNT signaling pathway in each genotype, which in CDX2-Het and CDX2-KO revealed conserved signaling between WNT5A and various frizzled receptors but near eliminated signaling between WNT5B and those same receptors, suggesting a critical role for WNT5B in the maintenance of the ncWNT pathway (Fig. 5B-C). We next examined trends in the clusters acting as senders, receivers, mediators, and influencers of the ncWNT pathway. In this analysis, mediators specifically control cell communication between any two groups, and influencers influence information flow more generally (Jin *et al*., 2021). Most clearly, we identified a marked decrease in the ability of CDX2-Het and CDX2-KO to act as senders and mediators in the three ExeM clusters (C1-C3). This pattern was also observed in PGCLCs, which share an extraembryonic origin (Sasaki *et al*., 2016).

**Figure 5:**
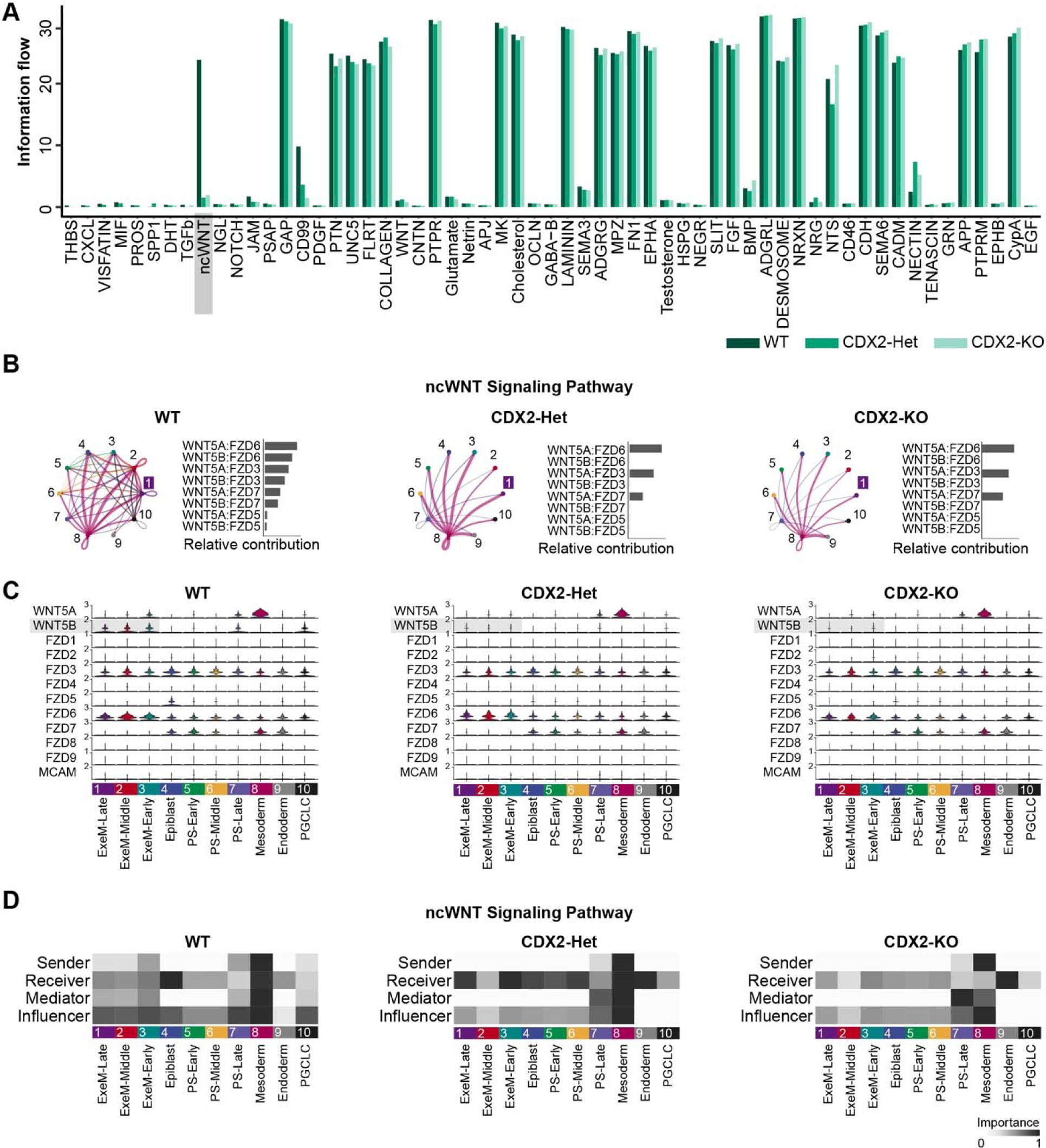
Cellchat reveals the CDX2-dose-dependent regulation of ncWNT signaling pathway. (**A**) Information flow (arbitrary units; A. U.) of various pathways across all clusters within WT, CDX2-Het, or CDX2-KO populations. (**B**) Circle plots (left) and bar plots (right) visualizing ligand-receptor communication across all clusters. The number correlates with cluster identity, and “1” indicates ExeM-Late. Line thickness corresponds with the strength of the predicted communication. (**C**) Violin plot of gene expression for each ligand or receptor comprising the ncWNT signaling pathway. (**D**) Heatmap showing the communication dynamics for the ncWNT signaling pathway in each cluster.

This pattern, in conjunction with the snRNA-seq data, suggests that the changes in information flow across genotypes are largely due to the inability of CDX2-Het and CDX2-KO to express WT levels of *WNT5B* in their extraembryonic populations, and implicate the non-canonical WNT signaling pathway in regulating early extraembryonic mesoderm development in a CDX2 dose-dependent manner. WNT5B has been shown to influence both canonical WNT/ β-catenin signaling and VEGF-C expression (Kanazawa *et al*., 2005), both of which have been shown to regulate vasculogenesis *in vivo* (Cao *et al*., 1998; Drake *et al*., 2000; Kanazawa *et al*., 2005; Shibuya, 2011). Additionally, components of the ncWNT pathway, including WNT5 and WNT11, have been shown to influence angiogenesis via regulation of the VEGF inhibitor FLT1 (Stefater *et al*., 2011; Akoumianakis, Polkinghorne and Antoniades, 2022). Thus, CDX2-driven *WNT5B* expression may be required to properly regulate these two pathways and sustain development within the allantois. Our current ongoing efforts have so far not yielded definitive results but so far do not negate this possibility.

### CDX2 and TBXT jointly regulate genes involved in extraembryonic mesoderm development

Because CDX2 and TBXT null animals share an embryonic lethal defect in allantois development and chorioallantoic fusion, we next asked whether we could identify common genes that are misregulated during extraembryonic mesoderm development both CDX2-KO and TBXT-KO conditions. We recently conducted a parallel analysis of a TBXT allelic series in the 2D gastruloid model that uses the same WT cell population as this current study (Bulger *et al*., 2023). We isolated the analogous extraembryonic mesoderm population from this TBXT dataset (“Extraembryonic-Late”), conducted TBXT-Het vs. WT and TBXT-KO vs. WT comparisons within this cluster, and looked at how the resulting lists of differentially expressed genes intersected with those of the CDX2 allelic series ExeM-Late population.

From this comparative analysis, we uncovered 11 genes downregulated and 32 genes upregulated in TBXT-Het vs. WT and 35 downregulated and 49 upregulated genes in TBXT-KO relative to WT (Fig. 6A, C, Table S4). Of the 35 genes downregulated in TBXT-KO, 8 were also downregulated in CDX2-KO relative to WT. These 8 genes included *ANK3, LSAMP,* and *ANGPT1*, which regulate adhesions and the organization of the actin cytoskeleton (Babcock *et al*., 2018; Liu *et al*., 2021), and the canonical WNT inhibitor *DCDC2* (Fig. 6B). 5 of these 8 genes were also significantly downregulated in CDX2-Het and 0 were significantly downregulated in TBXT-Het, although trends toward intermediate expression are evident (p-adj < 0.05, Log2FC > 0.25) (Fig. 6B, Table S4). Notably, both *ANK3* and *ANGPT1* have been shown to regulate angiogenesis *in vivo* via the VEGF signaling pathway (Gavard, Patel and Gutkind, 2008; Cao *et al*., 2017; Liu *et al*., 2021). VEGF increases vascular permeability while ANGPT1 and ANK3 reduce permeability (Senger *et al*., 1983; Thurston *et al*., 1999; Liu *et al*., 2021), and the proper balance of these factors is likely required for proper placental vasculogenesis. In addition, both ANGPT1 and ANK3 reduce the cell surface localization of ß-catenin and endothelial barrier function. This function is impaired in ANGPT1 heterozygotes (Durak *et al*., 2015; d’Apolito *et al*., 2019), suggesting that proper regulation of their expression is required for the development of extraembryonic-mesoderm-derived structures.

**Figure 6.**
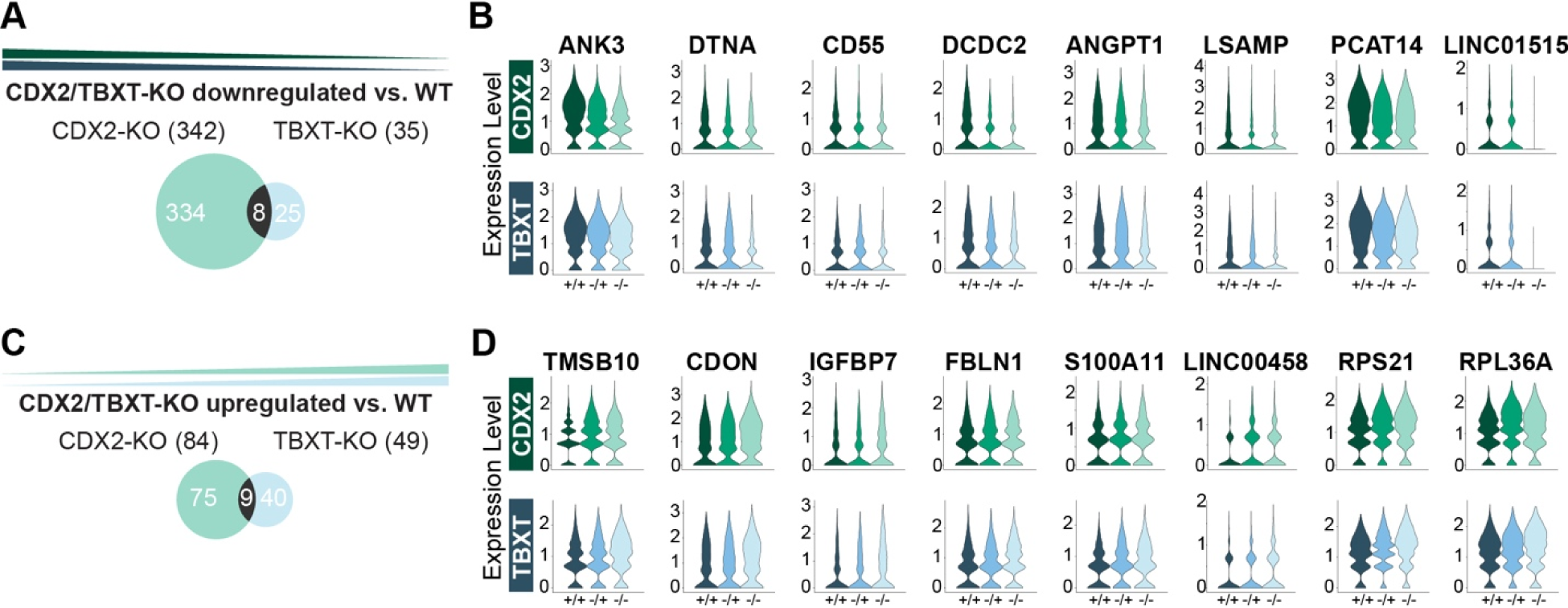
TBXT and CDX2 mutants both misregulate genes associated with the VEGF signaling pathway. (**A**) Venn diagram illustrating overlapping genes downregulated or (**C**) upregulated in CDX2-KO (left) or TBXT-KO (right) relative to WT in the ExeM-Late cluster. (**B**) Violin plots of gene expression for overlapping genes identified in (**A**). MT-ND3 shown in Fig. S9. (**D**) Violin plots of gene expression for overlapping genes identified in (**C**).

Of the 49 upregulated genes in TBXT-KO, 9 were also upregulated in CDX2-KO (Fig. 6C). These 9 genes included the cytoskeleton modulator *TMSB10*, hedgehog pathway and myogenesis effector *CDON*, and placental adhesion regulators *FBLN1* and *IGFBP7* (Fig. 6D, Table S4). Like *ANK3* and *ANGPT1*, *TMSB10* influences VEGF expression, specifically inhibiting VEGF-induced endothelial cell proliferation, migration, and invasion (Lee *et al*., 2005; Pan *et al*., 2020). Five of these 9 genes were also significantly upregulated in CDX2-Het and 0 were upregulated in TBXT-Het (Table S4). These results suggest that one method by which CDX2 and TBXT regulate the development of extraembryonic mesoderm structures is by promoting the expression of *ANGPT1* and *ANK3*, which in turn sequester β-catenin in the cytoskeleton, preventing its nuclear translocation and fine-tuning the expression of downstream canonical WNT effectors. These factors also likely influence the activity of the VEGF signaling pathway and the associated development of extraembryonic mesoderm.

Taken together, the changes in gene expression shared between the TBXT-KO and CDX2-KO compared to WT reveal effectors that regulate cytoskeletal architecture, cell adhesions, and cell permeability via the VEGF and WNT signaling pathways. The canonical WNT pathway regulates several pro-angiogenic molecules including VEGF family members, whose expression in turn positively correlates with cytoplasmic β-catenin localization (Kasprzak, 2020). *In vivo,* proper regulation of these factors is likely required for the development of the allantois and subsequent chorioallantoic fusion, and their misregulation contributes to the early embryonic lethality observed *in vivo* in the absence of TBXT or CDX2.

## DISCUSSION

Proper morphogenesis of extraembryonic structures is critical for an embryo to develop to term. CDX2 specifically is required for the development of several extraembryonic-mesoderm-derived structures, including the growth of the allantois, chorioallantoic fusion, and yolk sac vasculogenesis, and its absence leads to early embryonic lethality. In this study, we sought to understand the gene regulatory network underlying the development of these structures and how it is dysregulated in the absence of CDX2. We additionally asked the extent to which this network is affected in a dose-dependent manner, motivated by studies showing both a correlation between CDX2 expression levels and the development of embryonic mesoderm, including elongation of the tailbud and studies showing that genes in related pathways such as TBXT can dose-dependently regulate allantois development.

Through this work, we demonstrate that relative to WT, both CDX2-Het and CDX2-KO have altered expression of both canonical and non-canonical WNT and BMP signaling pathways, in addition to changes in cell adhesions and cytoskeletal regulators. Over 2/3 of the genes misregulated in CDX2-Het are also misregulated in CDX2-KO, revealing a striking dose dependence in gene expression downstream of CDX2. Is it therefore likely that the changes in gene expression observed in CDX2-Het correlate with the impaired or delayed development of the allantois and associated vasculature observed *in vivo*. However, this impairment is likely not sufficient to prevent chorioallantoic fusion and cause lethality, as evidenced by the viability of heterozygous CDX2 mice. This prediction agrees with the phenotype observed in mice heterozygous for TBXT.

Even with these changes in gene expression, we observe far fewer differentially accessible peaks in CDX2-Het than in CDX2-KO when compared to WT. The peaks identified in CDX2-KO are heavily enriched for CDX motifs, suggesting that CDX2 binding drives their accessibility. Additionally, we observe that CDX2-Het binds to CDX motifs at a reduced frequency compared to WT, but CDX2-Het and WT share very similar chromatin accessibility profiles. This observation suggests that reduced CDX2 expression is sufficient to drive a WT-like chromatin accessibility profile, even if downstream gene expression is compromised. This perhaps indicates that open chromatin is not sufficient to drive the expression of downstream genes and that a threshold amount of CDX2 must be bound to regulatory elements for expression to reach WT levels. It is also possible that a balance between CDX2 and its co-factors is required to activate downstream gene expression, and this balance is not achieved in CDX2-Het despite the chromatin being accessible.

We next demonstrate that CDX2 dose influences HOX gene expression patterns and identify HOXB paralog as being uniquely regulated in our dataset. The detection of the HOXB cluster perhaps reflects a temporal delay in expression between different HOX paralogs, or it may be a reflection of a HOXB-specific role in the extraembryonic mesoderm. Regardless, we observe a reduction in central HOX gene expression in WT relative to both CDX2-Het and CDX2-KO, again emphasizing how CDX2 dose influences the expression of its downstream targets. Additionally, we observe the upregulation of *HOXB2* in CDX2-KO, reminiscent of the anterior homeotic shifts seen *in vivo*.

We next look at how communication between ligands and receptors of various clusters is influenced by CDX2 dose using the R package, CellChat. Through this analysis, we isolate the ncWNT pathway as uniquely misregulated in both CDX2-Het and CDX2-KO relative to WT. Changes in the ncWNT signaling pathway are largely restricted to signals from and within the three ExeM clusters and specifically reflect the reduction in *WNT5B* expression in the ExeM of the mutant cell lines. *WNT5B* influences both canonical WNT signaling and VEGF signaling, both established regulators of early embryonic vasculogenesis in vivo, thus implicating WNT5B expression as critical for early morphogenesis of extraembryonic mesoderm-derived structures.

Finally, we compare how the CDX2-KO and TBXT-KO influence extraembryonic mesoderm gene expression and isolate genes that are shared between the two datasets. These genes are reflective of adhesions and cytoskeletal dynamics, and several converge on the VEGF signaling pathway, suggesting that CDX2 and TBXT both disrupt pathways that have been shown *in vivo* to be critical for early vasculogenesis. These pathways likely contribute to malformations in the development of the allantois, preventing chorioallantoic fusion and subsequent placental development.

Taken together, these results clarify the dose-dependent role of CDX2 in the formation of extraembryonic structures crucial for early embryogenesis. Understanding the genetic patterns underlying the development of these structures is critical for our foundational understanding of how gene dosage influences morphogenesis, chromatin conformation, and downstream gene expression, with the ultimate goal of better understanding extraembryonic mesoderm and subsequent placental development.

## METHODS

### Cell Lines

All work with human induced pluripotent stem cells (hiPSCs) was approved by the University of California, San Francisco Human Gamete, Embryo, and Stem Cell Research (GESCR) Committee. Human iPS cells harboring genome-edited indel mutations for *CDX2* (*CDX2^+/-^ (CDX2-Het) and CDX2^-/-^ (CDX2-KO*)) were generated for this study and derived from the Allen Institute WTC11-LaminB parental cell line (AICS-0013 cl.210). The WT line was derived from a WTC11-LaminB subclone that was exposed to a TBXT sgRNA (CAGAGCGCGAACTGCGCGTG; a gift from Jacob Hanna (Addgene plasmid #5972)) but remained unedited. All cell lines were karyotyped by Cell Line Genetics and reported to be karyotypically normal. Additionally, all cell lines tested negative for mycoplasma using a MycoAlert Mycoplasma Detection Kit (Lonza).

### Maintenance of iPS Cells

Human iPSCs were cultured on growth factor-reduced (GFR) Matrigel (Corning Life Sciences) and fed at minimum every other day with mTeSR-Plus medium (STEMCELL Technologies) (Ludwig et al., 2006). Cells were passaged by dissociation with Accutase (STEMCELL Technologies) and re-seeded in mTeSR-Plus medium supplemented with the small molecule Rho-associated coiled-coil kinase (ROCK) inhibitor Y276932 (10 μM; Selleckchem) (Park et al., 2015) at a seeding density of 12,000 cells per cm^2^. After 24 hours, cells were maintained in mTeSR-Plus media until 80% confluent.

### CDX2 Allelic Series Generation

To generate the CDX2 allelic series we first lipofected WTC11-LaminB cells (AICS-0013 cl.210) with 125ng sgRNA and 500ng Cas9 protein according to the Lipofectamine Stem Transfection Reagent Protocol (Invitrogen). Two CDX2 sgRNAs that both target the first exon of CDX2 gene were co-lipofected using 62.5ng each: (CDX2-A: CCUAGCUCCGUGCGCCACUC and CDX2-B: AGUUCUGCGGCGCCAGGUUG). After recovery for 48 hours in mTeSR-Plus supplemented with ROCK inhibitor, lipofected cells were dissociated using Accutase and passaged into a GFR-Matrigel coated 10cm dish, where they were expanded for 24hr in mTeSR-Plus with ROCK inhibitor. Media was replaced with mTeSR-Plus without ROCK inhibitor and cells continued to grow another 2-4 days before the manual selection of 20-30 single colonies into individual wells of a 96-well plate. After the expansion of the clonal populations for 5-10 days, cells were passaged into a new 96-well plate at a 1:5 dilution ratio in mTeSR-Plus supplemented with ROCK inhibitor, and the remaining cells were used for genotyping. For screening of *CDX2* exon 1 non-homologous end-joining (NHEJ) mutations, DNA was isolated using QuickExtract DNA lysis solution (Epicentre #QE0905T), and genomic DNA flanking the targeted sequence was amplified by PCR (For1: cctcgacgtctccaaccattg and Rev1: gcctctgcttaccttggctg) and sequenced using Rev1. Synthego ICE analysis was employed to quantify editing efficiency and identify clones with heterozygous (45-55% KO score) or homozygous null (>90% KO score) mutations (Table S1). To eliminate the possibility of a heterozygous line being a mixed population of wildtype and homozygous null alleles, 8 subclones of the prospective heterozygous line were isolated and Sanger sequenced as before, and all subclones were confirmed to contain identical genotypes (Table S1). After sequencing confirmation of respective genotypes, karyotypically normal cells from each hiPSC line (Fig. S1) were expanded for subsequent studies.

### PDMS stamp fabrication

Stamps were fabricated as described in Bulger et al. 2023. Standard photolithography methods were used to fabricate a master template, which was provided as a gift from PengFei Zhang and the Abate lab at the University of California, San Francisco (Alom Ruiz and Chen, 2007; Théry and Piel, 2009; Minn *et al*., 2020). The photoresist master was then coated with a layer of chlorotrimethylsilane in the vacuum for 30 min. Polydimethylsiloxane (PDMS) and its curing agent, Sylgard 184, (Dow Corning, Midland, MI) were mixed in a 10:1 ratio, degassed, poured over the top of the master, and cured at 60°C overnight, after which the PDMS layer was peeled off to be used as a stamp in micro-contact printing.

### Microcontact Printing

Matrigel stamps were applied as described in Bulger et al. 2023. PDMS stamps (each containing 12 x 1000uM circles) were sterilized by washing in a 70% ethanol solution and dried in a laminar flow hood. Growth factor reduced Matrigel (Corning) was diluted in DMEM/F-12 (Gibco) at 1:100 dilution and incubated on the stamps to cover the entire surface of the feature side at 37°C for 1 hour. The Matrigel solution was then aspirated off the stamps, which were air-dried. Using tweezers, the Matrigel-coated surface of stamps was brought in contact with glass or plastic substrate, usually a glass 24-well plate or removable 3-chamber slide (Ibidi), and incubated on the substrate for 1hr at 37°C. The stamps were then removed and rinsed in ethanol for future use. Matrigel-printed substrates were incubated with 1% Bovine Serum Albumin (Sigma Aldrich) in DPBS-/- at room temperature for 1 hour before being stored in DPBS -/- solution at 4°C for up to 2 weeks.

### Confined 2D Gastruloid Differentiation

2D gastruloids were differentiated as described in Bulger et al. 2023. hiPSCs were dissociated with Accutase and resuspended in mTeSR-Plus supplemented with ROCK inhibitor. Cells were then seeded onto a stamped well at a concentration of approximately 750 cells/mm^2^. Cells were incubated at 37°C for 3 hours before the well was rinsed 1x with DPBS and given fresh mTeSR-Plus supplemented with ROCK inhibitor. Approximately 24 hours post-seeding, media was exchanged for mTeSR-Plus. After another 24 hours or upon confluency of the stamped colony, media was exchanged for mTeSR-Plus supplemented with BMP4 (50ng/mL). Colonies were allowed to differentiate in the presence of BMP4 for 48 hours before being processed for downstream analyses.

### Western Blot

Cells of each genotype were induced to form CDX2+ extraembryonic mesoderm in a sparsely seeded monolayer exposed to 50ng/uL BMP4 in mTeSR+ for 48 hours before protein isolation. Cells were washed twice with ice-cold PBS and lysed in RIPA lysis buffer (Fisher Scientific; A32965). Three replicate wells were pooled for each genotype for each differentiation condition. The protein concentration was determined using the Pierce BCA Protein Assay Kit (Life Technologies, 23227) and quantified on a SpectraMax i3 Multi-Mode Platform (Molecular Devices) following the manufacturer’s instructions. Protein (∼20-40 μg) was transferred to the membrane using the Trans-Blot Turbo Transfer System (Biorad; 1704157). The membrane was then blocked overnight at 4°C using Intercept TBS Blocking Buffer (Li-COR; 927-70001). Primary antibodies CDX2 (12306; 1:1000; Rb), ß-Actin (ab8226; 1:1000; Ms), or α-Tubulin (T5168; 1:1000; Ms) were diluted in Intercept T20 (TBS) Antibody dilution buffer (Li-COR) at a 1:1000 ratio and incubated with the membrane overnight at 4°C. The next morning, membranes were washed in 1x TBS-T and incubated for 1 hour at RT in the dark with species-specific secondary antibodies (Rb-680; 926-68071; Ms-800 926-32212) (VWR) at 1:10,000. Membranes were subsequently washed and developed using the BioRad ChemiDoc MP. Protein levels were quantified using ImageJ by first subtracting the intensity of a blank ROI from the experimental ROI, and then calculating a normalization factor by dividing the observed housekeeping intensity by the highest observed housekeeping intensity. The observed experimental signal was then divided by the lane normalization factor to generate a normalized experimental signal. Each lane from the same blot was then converted to a percentage of the highest WT normalized experimental signal on that blot.

### Immunofluorescence

Immunofluorescence was conducted as described in Bulger et al. 2023. hiPSCs were rinsed with PBS 1X, fixed in 4% paraformaldehyde (VWR) for 15-20 minutes, and subsequently washed 3X with PBS. The fixed cells were permeabilized and blocked in a buffer comprised of 0.3% Triton X-100 (Sigma Aldrich) and 5% normal donkey serum in PBS for one hour and then incubated with primary antibodies SOX2 (3579s; 1:200; Rb) CDX2 (ab157524; 1:500; Ms), or TBXT (AF2085; 1:400; Gt) diluted in antibody dilution buffer (0.3% Triton, 1% BSA in PBS) overnight. The following day, samples were washed 3X with PBS and incubated with secondary antibodies in antibody dilution buffer at room temperature for 2 hours. Secondary antibodies used were conjugated with Alexa 405, Alexa 555, or Alexa 647 (Life Technologies) at a dilution of 1:400. Cells were imaged at 10x, 20x, or 40x magnification on an inverted AxioObserver Z1 (Zeiss) with an ORCA-Flash4.0 digital CMOS camera (Hamamatsu).

### Cell Harvesting for Single Nuclei Multiome ATAC + RNA Sequencing

Each of the *CDX2* genotypes was differentiated, harvested, and prepared at the same time for each of the two biological replicates, as described in Bulger et. al. 2023. Therefore, each set of biological replicates represents an experimental batch. For each genotype (sample) within the batch, 12 micropatterns were differentiated within each well of a 24-well plate and cells from all wells on a plate were pooled, yielding a cell suspension comprising approximately 288 colonies per sample. Nuclei were isolated and ∼10,000-19,000 nuclei/sample were transposed and loaded onto a 10x Chromium Chip J to generate gel bead-in emulsions (GEMs) following the 10x Chromium Next GEM Single Cell Multiome ATAC and Gene Expression Kit (10x Genomics, CG000338). WT-1 (eb01)= 12,072 nuclei, 20,556.09 reads/nucleus. CDX2-KO (eb04) = 18,239 nuclei, 15,288.63 reads/nucleus. CDX2-Het-1 (eb05) = 18,806 nuclei, 19,435.86 reads/nuclei. WT-2 (eb06) = 10,903 nuclei, 26,003.63 reads/nucleus. CDX2-KO-2 (eb09) = 12,314 nuclei, 20,150.10 reads/nucleus. CDX2-Het-2 (eb10) = 11,602 nuclei, 18,929.24 reads/nucleus. GEMs were processed to produce ATAC and gene expression libraries in collaboration with the Gladstone Genomics Core. Deep sequencing was performed on the NovaSeq 6000 S4 200 cycle flow cell for a read depth of >15k reads per cell.

### Data Processing Using CellRanger-Arc

All ATAC and GEX datasets were processed using CellRanger-Arc 2.0.0. FASTQ files were generated using the mkfastq function, and reads were aligned to the hg38 reference genome (version 2.0.0).

### Seurat Analysis

Outputs from the CellRanger-Arc count pipeline were analyzed using the Seurat package (version 4.3.0)(Satija *et al*., 2015; Butler *et al*., 2018; Stuart *et al*., 2019) in R (v4.2.0). Quality control filtering included the removal of outliers due to the number of features/genes (nFeature_RNA > 2500 & nFeature_RNA < 4500, nCount_RNA > 200 & nCount_RNA < 12000, mitochondrial percentage > 5% and mitochondrial percentage < 20%, and ribosomal percentage > 3% and ribosomal percentage < 10%). Cell cycle scores were added using the function CellCycleScoring. ScTransform v2 normalization was then performed to integrate samples based on batch with regression based on cell cycle scores and ribosomal content (vars.to.regress = c(“S.Score”, “G2M.Score”, “percent_ribo”)). Principal component analysis (PCA) was performed using the most highly variable genes, and cells were clustered based on the top 15 principal components using the functions RunUMAP, FindNeighbors, and FindClusters, and the output UMAP graphs were generated by DimPlot. The resolution parameter of 0.4 was set so that cluster boundaries largely separated the likely major cell types. Cluster annotation was performed based on the expression of known marker genes, leading to 10 broadly assigned cell types. Cells filtered out of the ArchR dataset based on doublet identification (see” ArchR Analysis” below) were removed from the Seurat dataset (final n = 25,557 cells). Differential gene expression was then performed with the function FindAllMarkers (logfc.threshold = 0.25 and min.pct = 0.1) to generate a list of top marker genes for each cluster. In pairwise comparisons of differential gene expression, positive values reflect upregulation in mutant lines, while negative values reflect upregulation in WT.

### ArchR Analysis

Indexed Fragment files generated by the CellRanger-Arc counts function served as input for the generation of sample-specific ArrowFiles (minTSS = 4 & minFrags = 1000) using the R package ArchR v1.0.2(Granja *et al*., 2021). ArrowFile creation also generates a genome wide TileMatrix using 500bp bins and a GeneScoreMatrix, an estimated value of gene expression based on a weighted calculation of accessibility within a gene body and surrounding locus. Each Arrow file (n=6 total) was then aggregated into a single ArchRProject for downstream analysis. Corresponding Gene Expression Matrices were imported to the project based on the filtered feature barcode matrix h5 file generated by CellRanger-arc counts and descriptive cluster labels were imported from the corresponding Seurat object based on cell barcodes. Cells filtered out of the Seurat dataset based on QC metrics previously described were also removed from the ArchR dataset. Cell doublet removal was performed in ArchR using the functions addDoubletScores and filterDoublets, leaving 25,557 cells with a median TSS of 13.431 and a median value of 11,458 fragments per cell (cells filtered = WT-1 0/2908, CDX2-Het-1 257/5072, CDX2-KO-1 0/ 2836, WT-2 288/5368, CDX2-Het-2 = 243/4935, CDX2-KO-2 = 306/5532).

After generation of the aggregated ArchR project, dimensionality reduction was performed using ArchR’s implementation of Iterative Latent Semantic Indexing (LSI) with the function addIterativeLSI based on the 500bp TileMatrix with default settings (iterations = 2, sampleCells = 10000, n.start = 10, resolution = 2, maxClusters = 6). This was repeated using the Gene Expression Matrix based on 2,500 variable features, yielding “LSI-ATAC” and LSI-RNA” reduced dimensions, respectively. The two reduced dimension values were then combined using addCombinedDims to yield “LSI_Combined,” which was used as input for batch correction using Harmony with the function addHarmony (groupby = “Sample, “Batch”). Clustering was then performed using Harmony-corrected values with addClusters with a resolution of 0.4 from the R package Seurat. Finally, clusters were visualized with function plot embedding, using batch-corrected single-cell embedding values from Uniform Manifold Approximation and Projection (UMAP) using the function addUMAP. Clusters and their corresponding UMAP projection were very similar to those generated based on RNA data in Seurat, and unless otherwise stated cluster identities in figures are based on barcodes transferred from Seurat rather than ArchR’s LSI implementation.

After cluster annotation, pseudobulk replicates of cells within similar groups were created to facilitate peak calling. Replicates were created using addGroupCoverages and peak calling was performed using addReproduciblePeakSet using standard settings by implementing MACS2. We then used ArchR’s iterative overlap peak merging method to create a union peakset of 305,429 unique peaks.

Cluster-enriched marker peaks were identified with getMarkerFeatures, using a Wilcoxon test and normalizing for biases from TSS enrichment scores and sequencing depth, and visualized with plotMarkerHeatmap, filtering for FDR =< 0.01 and abs(Log2FC) => 1.25. Motif enrichment of cluster-enriched peaks was done using addMotifAnnotations with the “cisbp” motif set. Enriched motifs per cluster were visualized by first running peakAnnoEnrichment, with FDR <= 0.1 and Log2FC >= 0.5. The top 20 significantly enriched motifs per cluster were visualized as a heatmap using plotEnrichHeatmap.

Peak-to-gene linkage analysis was performed in ArchR using the addPeak2GeneLinks command, using the batch-corrected Harmony embedding values. A total of 24,487 linkages were found using FDR 1e-04, corCutOff = 0.45, and a resolution of 1.

Differential accessibility within the extraembryonic mesoderm cluster was performed by using the command subsetArchrProject to subset the ArchR project based on the ‘ExeM-Late’ annotated cluster as determined from Seurat. This subsetting yielded 3,792 cells, with a median TSS of 13.396 and a median number of fragments of 11,419. Differentially expressed genes predicted pairwise across genotypes (WT vs. CDX2-KO or WT vs. CDX2-Het) were identified with getMarkerFeatures based on the GeneScoreMatrix, using a Wilcoxon Test and normalizing for biases from TSS enrichment scores and sequencing depth. GetMarkers was then run and visualized as a volcano plot using plotMarkers (FDR =< 0.1 and abs(Log2FC) >= 0.5). This process was repeated for the PeakMatrix to determine uniquely accessible peaks. 25 Peaks were detected between WT and CDX2-Het and 261 peaks were detected between WT and CDX2-KO. Significant ‘cisbp’ motif enrichments detected between genotypes within these peaks were calculated using peakAnnoEnrichment() (FDR <= 0.1 and abs(Log2FC) >= 0.5).

### Gene Ontology Analysis

Gene Ontology (GO) analysis for downregulated or upregulated CDX2-dependent genes was performed with ShinyGO V0.77 (Ge, Jung and Yao, 2020) using GO Biological Process or Molecular Function terms (FDR < 0.05, pathways size 2-2000). Downregulated or upregulated CDX2-dependent gene lists from the ExeM-Late subcluster were assembled from differential tests between *CDX2-Het vs. WT* or *CDX2-KO vs. WT* in Seurat. Gene sets were filtered with a significance threshold set at an adjusted p-value> 0.05 and the abs(log2FC) > 0.25. The first twenty hits for the CDX2-KO or CDX2-Het vs. WT comparison were visualized with lollipop plots. The process was repeated for genes identified as overlapping in both TBXT-KO and CDX2-KO vs. WT comparisons.

### CellChat

Cell signaling analysis was performed using the R package CellChat (Jin *et al*., 2021) as described in Bulger et al. 2023. The Seurat object containing all samples was subset by genotype, yielding a separate Seurat object for WT, CDX2-Het, or CDX2-KO. These 3 objects were then imported into CellChat using the function createCellChat. All ligand-receptor and signaling pathways within the CellChatDB.human were kept for analysis. Initial preprocessing to identify over-expressed ligands and receptors was performed using the functions identifyOverExpressedGenes and identifyOverExpressedInteractions with standard settings. Inference of cell communication was calculated with computeCommunProb(cellchat) and filtered by filterCommunication(cellchat, min.cells = 10). Pathway-level cell communication was calculated with computeCommunProbPathway, and aggregated networks were identified with aggregateNet, using standard settings. Network centrality scores were assigned with the function netAnalysis_computeCentrality. This workflow was run for WT, CDX2-Het, and CDX2-KO datasets independently and differential signaling analysis was then run by merging the WT, CDX2-Het, and CDX2-KO objects with mergeCellChat(). Information flow, which is defined by the sum of communication probability among all pairs of cell groups in the inferred network (i.e., the total weights in the network), was compared across genotypes using rankNet(cellchat). The distance of signaling networks between WT and TBXT-KO datasets was calculated by performing joint manifold learning and classification of communication networks based on functional similarity using computeNetSimilarityPairwise(cellchat), netEmbedding(cellchat), and netClustering (cellchat). Circle diagrams and heatmaps of pathways of interest were then generated for each genotype separately using the standard settings for netVisual_aggregate(cellchat) or netVisual_heatmap(cellchat), respectively. Violin plots of differential gene expression were generated using plotGeneExpression(cellchat) with the standard settings.

### Quantification and Statistical Analysis

Each experiment was performed with at least three biological replicates except multiomic snATAC and snRNA-seq, which was performed with two biological replicates. Multiple comparisons were used to compare multiple groups followed by unpaired *t*-tests (two-tailed) between two groups subject to a posthoc Bonferroni correction. In gene expression analysis, two replicates were used for each condition, and all gene expression was normalized to control wild-type populations followed by unpaired *t*-tests (two-tailed). Significance was specified as p-adj < 0.05 unless otherwise specified in figure legends. All error bars represent the standard error of the mean (s.e.m.) unless otherwise noted in the figure legend.

### Data and reagent availability

snATAC-seq and snRNA-seq data have been deposited in GEO under the accession number GSE245998 (WT) and GSE251813 (CDX2-Het and CDX2-KO). Analysis scripts used to generate figure panels and relevant cell lines are available from the authors upon request.

## ACKNOWLEDGEMENTS

We would like to thank all the members of the Bruneau and McDevitt labs for their continued support. We would like to thank Mylinh Bernardi and Horng-Ru Lin of the Gladstone Genomics Core, the Gladstone Histology and Light Microscopy Core, and the Gladstone Stem Cell Core for their experimental expertise. In addition, we thank Pengfei Zhang and the Abate Lab at UCSF for assistance with PDMS stamp fabrication.

## AUTHOR CONTRIBUTIONS

E.A.B., and T.C.M conceived and designed the study. E.A.B, performed all experimental work and data analysis of snATAC-seq and snRNA-seq data, including CellChat analysis. T.C.M. and B.G.B supervised and advised. E.A.B. prepared figures and wrote the manuscript with input from all co—authors.

---

**Figure S1:**
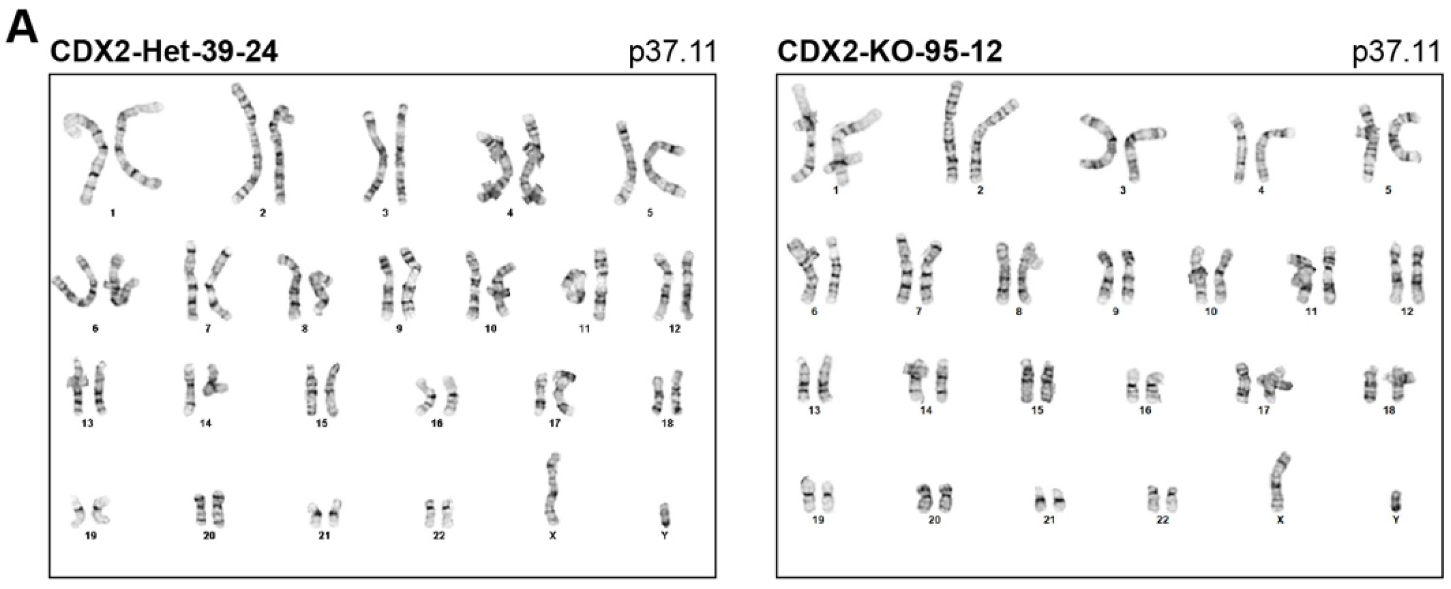
Karyotyping results from CDX2-Het and CDX2-KO isogenic lines. For WT, see Bulger et al. 2023.

**Figure S2:**
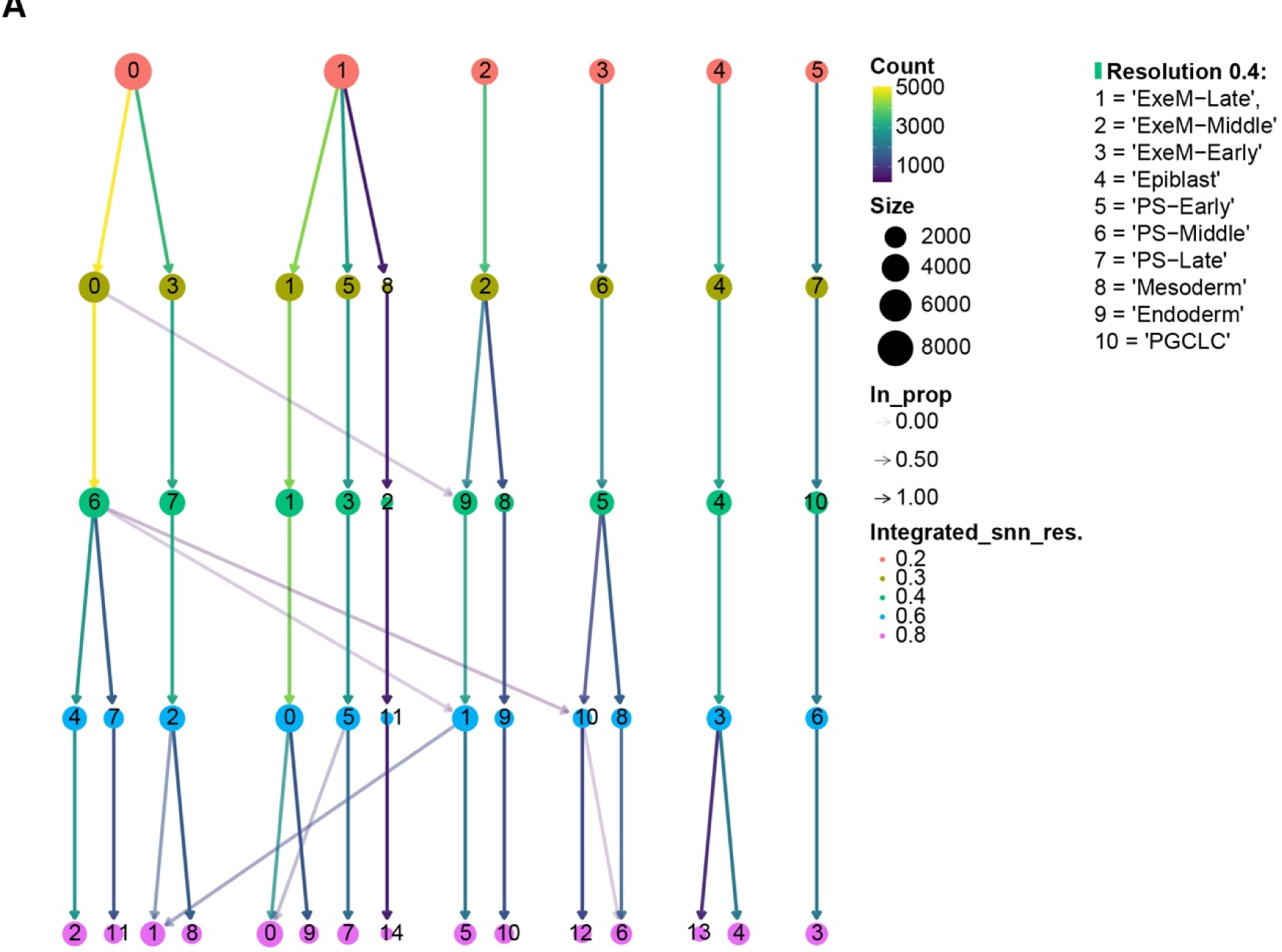
ClusTree analysis of clusters at 0.2, 0.3, 0.4, 0.6, and 0.8 resolution. A resolution of .4 was used for subsequent analyses.

**Figure S3:**
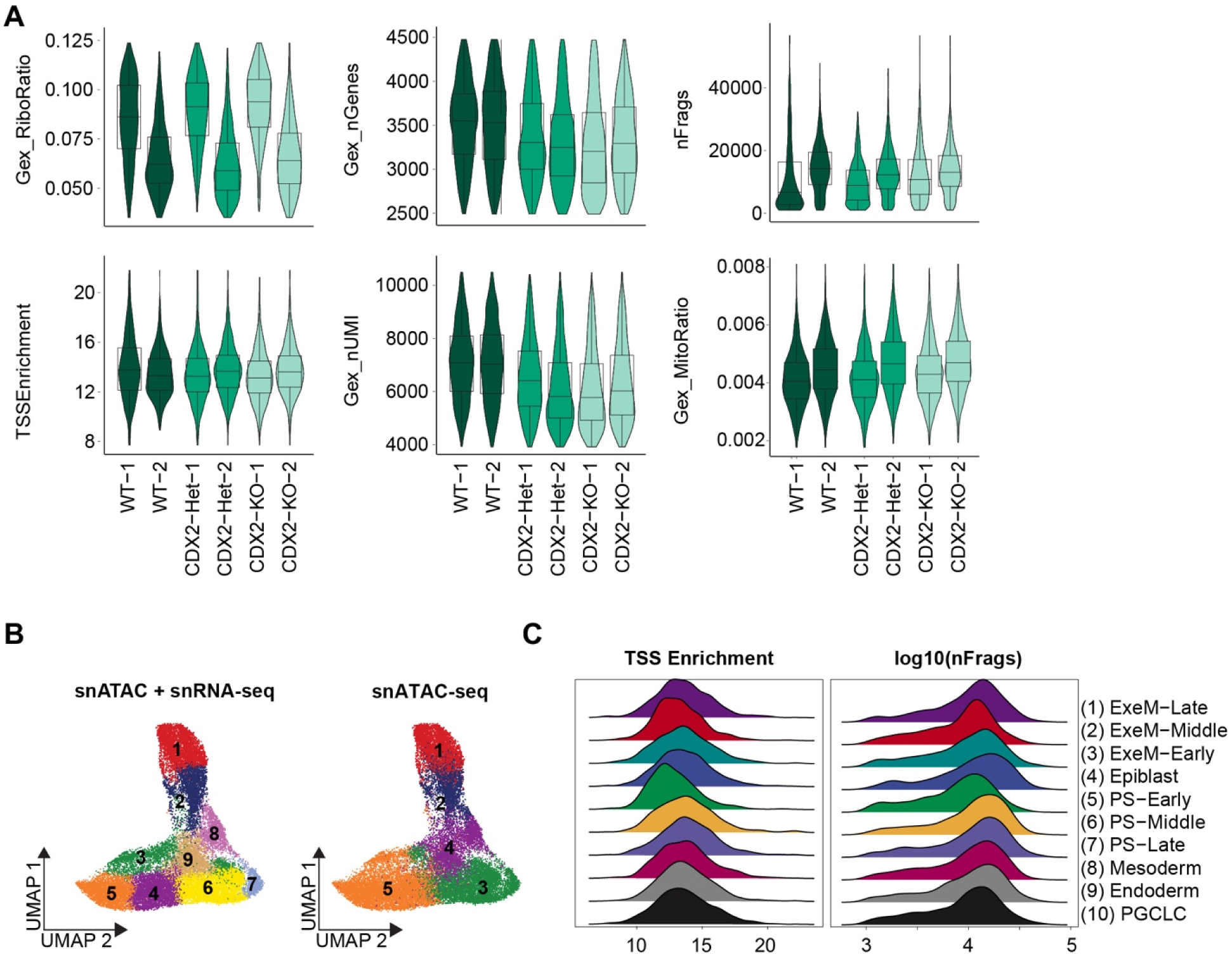
Quality control parameters for CDX2 snRNA-seq and snATAC-seq data. (**A**) Quality control parameters after filtration, separated by sample. (**B**) TSS enrichment and log10(nFrags), separated by cluster.

**Figure S4:**
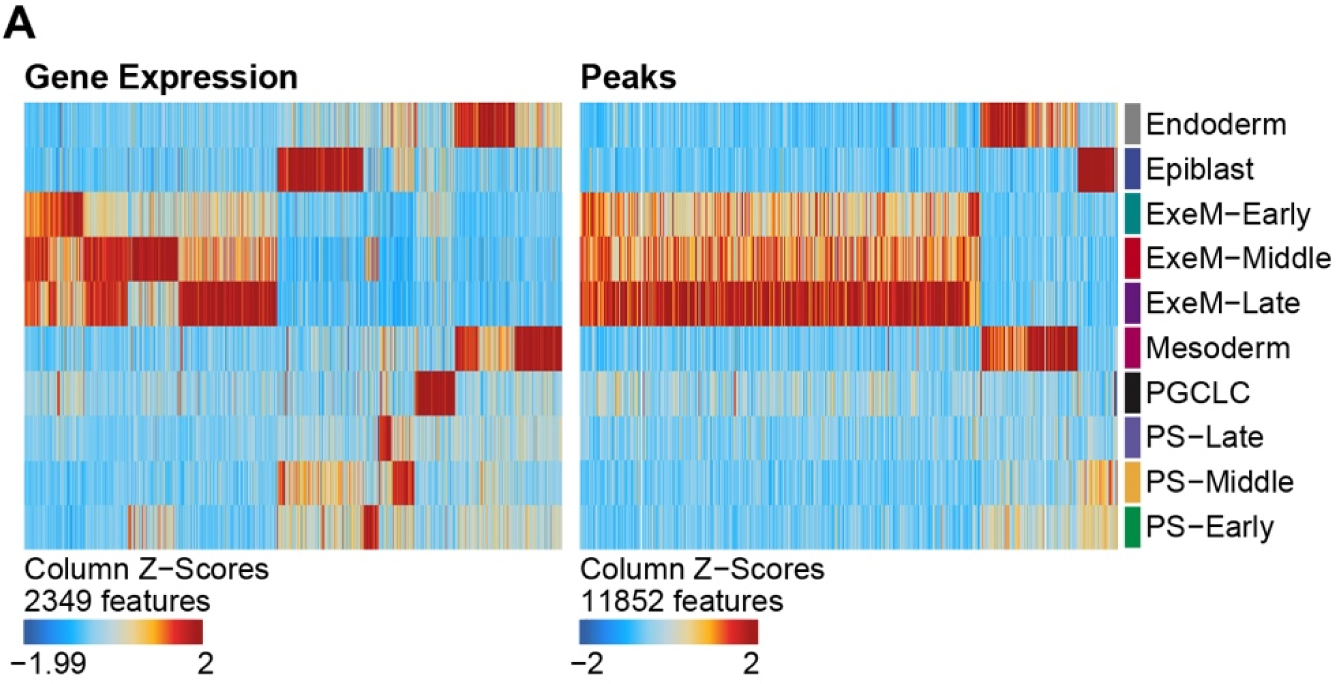
Gene expression and peak accessibility separated by cluster. The order of rows is based on hierarchical clustering.

**Figure S5:**
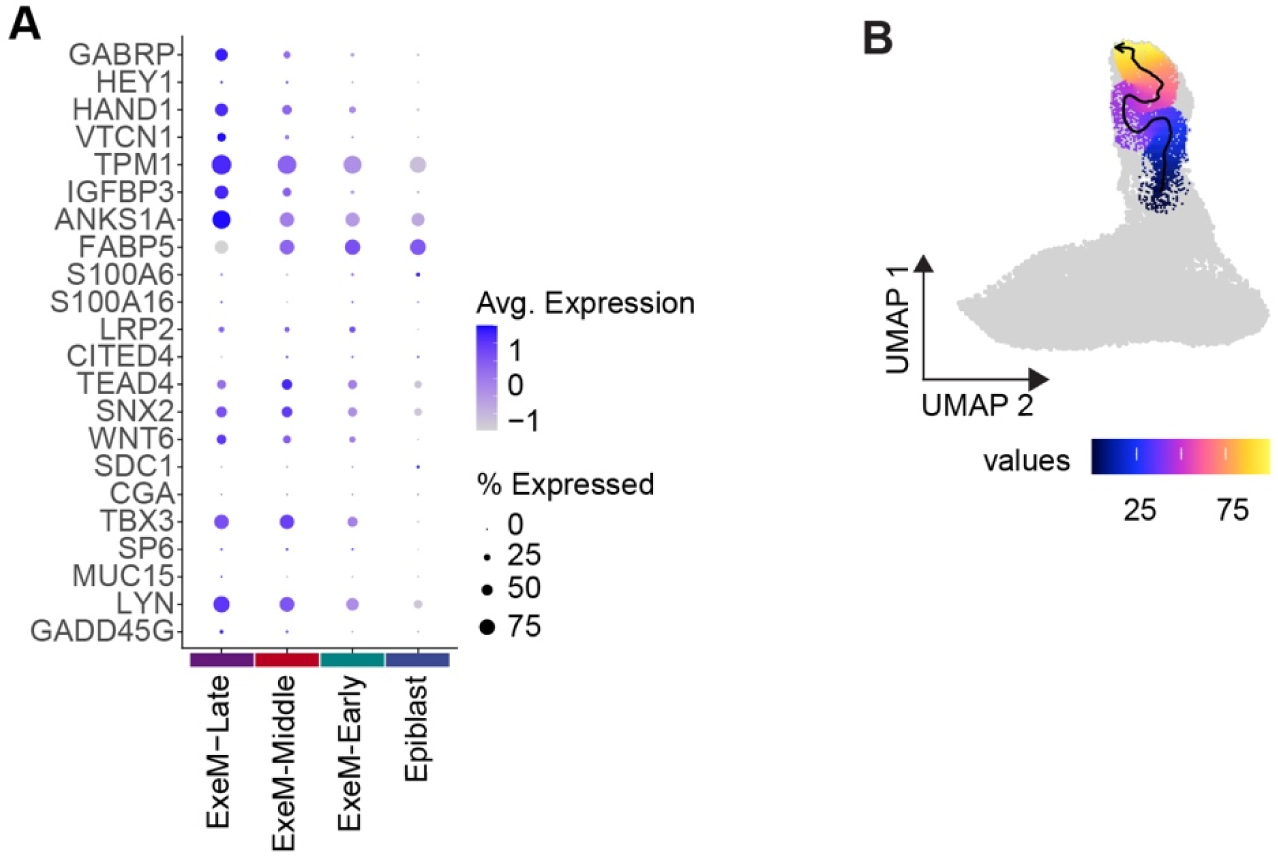
Extraembryonic mesoderm and trophectoderm marker expression across clusters. (**A**) Dotplot reflecting key markers of Amnion and trophectoderm for the extraembryonic and epiblast clusters (C1-C4). (**B**) Trajectory analysis of ExeM-Early, ExeM-Middle, and ExeM-Late (C3-C1).

**Figure S6:**
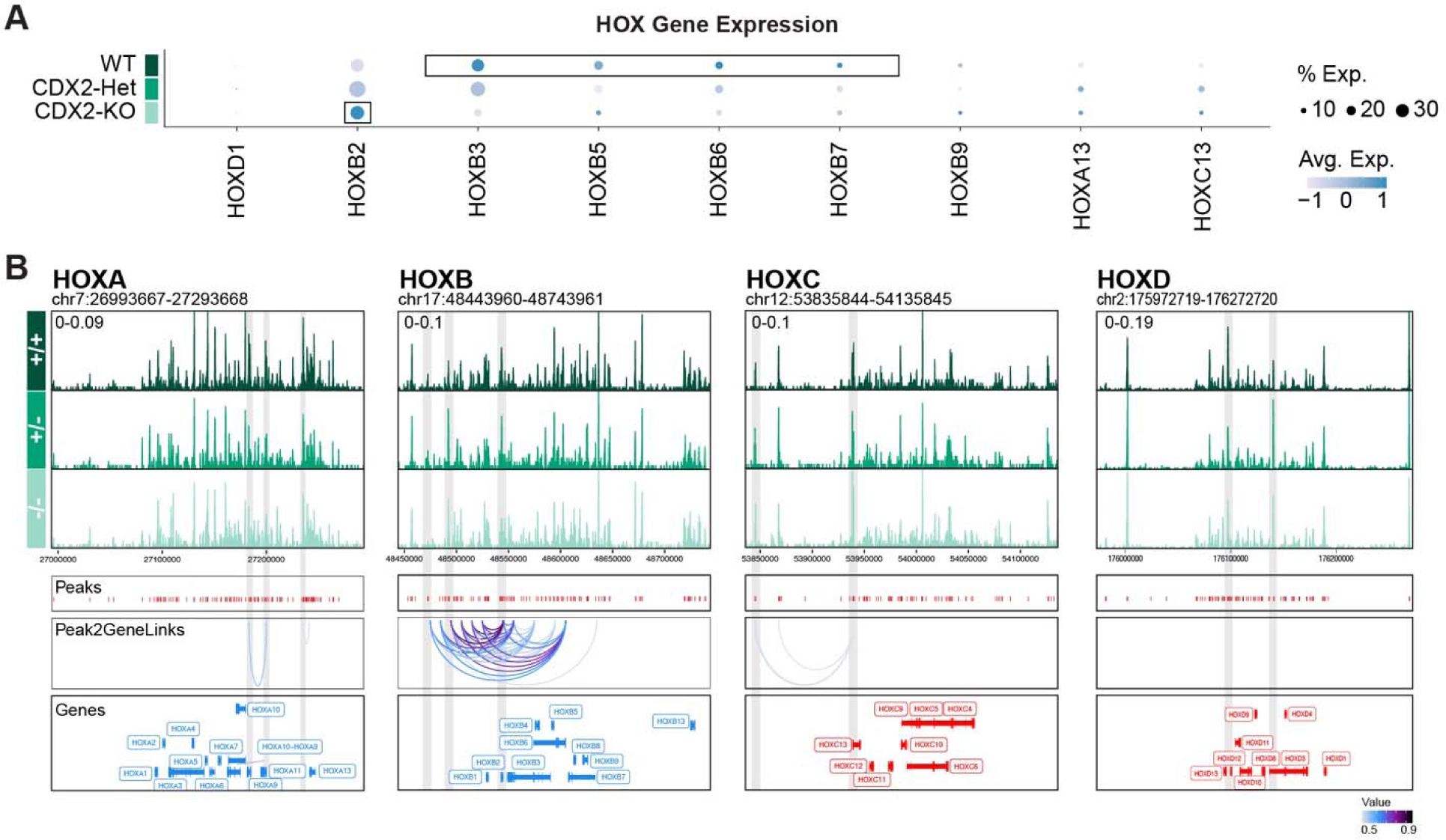
CDX2 dose-dependently influences downstream HOX expression. (**A**) Gene expression of detectable HOX genes in the EXEM-Late cluster across WT, CDX2-Het, and CDX2-KO. (**B**) Chromatin accessibility tracks in the EXEM-Late cluster across WT, CDX2-Het, and CDX2-KO with annotated peaks, peak2Gene links, and gene names.

**Figure S7:**
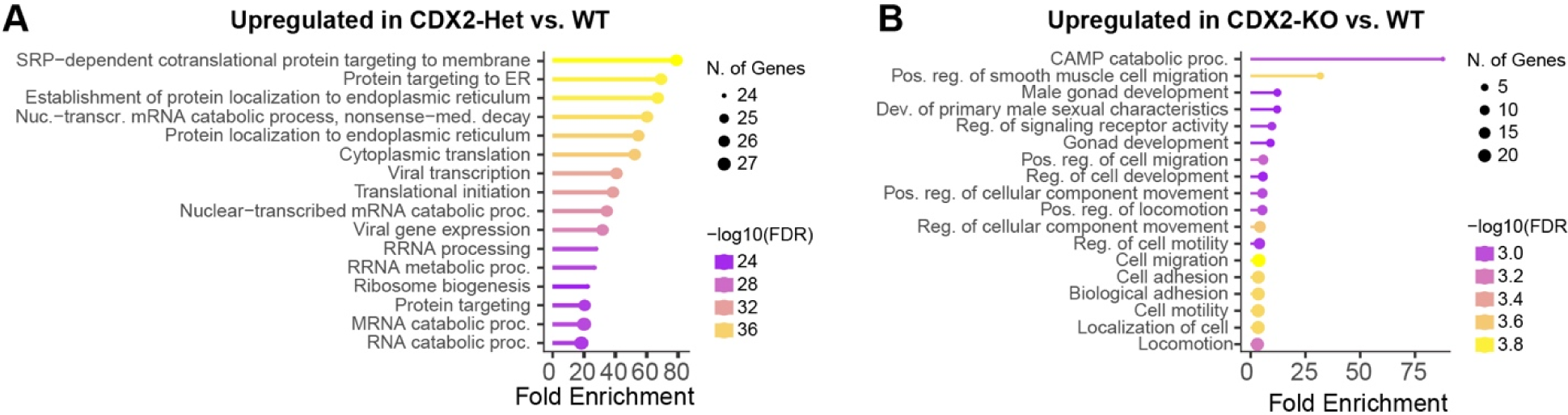
ShinyGO analysis of CDX2 allelic series. (**A**) ShinyGO analysis of genes upregulated in CDX2-Het vs. WT or (**B**) genes upregulated in CDX2-KO vs. WT (right) (Log2FC > −0.25, p-adj < 0.05).

**Figure S8:**
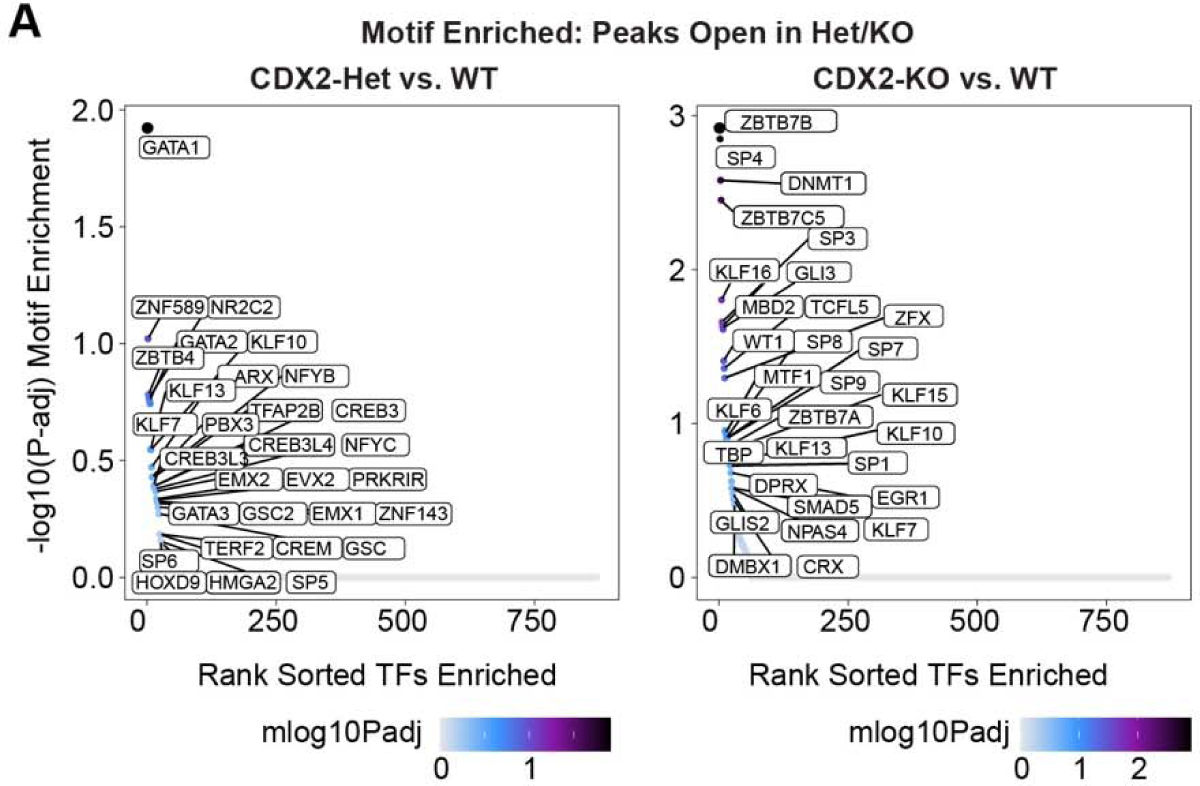
Motif enrichment in ExE-Late across CDX2 allelic series. (**A**) Motifs enriched in DARs more accessible in CDX2-Het relative to WT or (**B**) CDX2-KO relative to WT.

**Figure S9:**
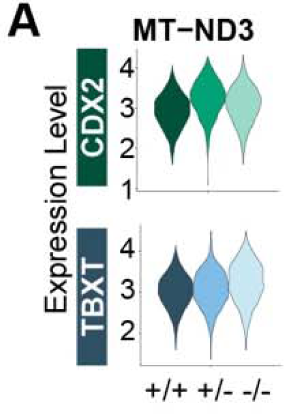
Gene Expression within the extraembryonic mesoderm cluster for MT-ND3 across CDX2 allelic series (top) or TBXT allelic series (bottom).

## SUPPLEMENTAL TABLES

**Table S1:** Indel frequency of clonal (#34, 39-24, 95-12) or subclonal (#39-24-2 through 39-24-12) cell populations exposed to the CDX2 sgRNA.

**Table S2:** Overview of differentially expressed genes (DEGs), differentially accessible regions (DARs), and differential peaks across genotypes within the extraembryonic mesoderm-late cluster.

**Table S3:** ShinyGO analysis.

**Table S4:** Shared differentially expressed genes (DEGs) in CDX2 and TBXT allelic series relative to WT.

